# Molecular basis of Salla Disease: R39C Mutation Effects on the Lysosomal Transporter Sialin

**DOI:** 10.64898/2026.04.20.719580

**Authors:** Christos Matsingos, Isaure Lot, Malaury Vaz, Jade Mailliart, Meryam Boulayat, Cécile Debacker, Anne Goupil-Lamy, Bruno Gasnier, Francine C. Acher, Christine Anne

## Abstract

Salla disease is caused by a genetic mutation in sialin, a lysosomal membrane transporter, which exports sialic acid from lysosomes. Substrate translocation occurs via a rocker-switch mechanism that alternately exposes the substrate-binding site to the lysosomal lumen and the cytosol. The pathogenic mutation R39C found in most Salla disease patients decreases the lysosomal localisation and the transport activity. In this study, we used computational and mutagenesis approaches to elucidate the molecular effects of the R39C mutation. Using three-dimensional models of human sialin in the lumen-open (LO) and cytosol-open (CO) states combined with the mutagenesis of selected residues, we identify a critical “triplet” motif comprising R39, E194, and E262, which is associated with an ionic lock formed between K197 and D350 in the LO conformation. Molecular dynamics simulations suggest that the electrostatic triplet negatively modulates the ionic lock, and are consistent with a strengthened ionic lock in R39C sialin, potentially favouring the LO state. To assess the global effects of the R39C mutation, we computed dynamic cross-correlation matrices and identified correlation patterns consistent with an allosteric coupling between the ionic lock K197/D350 and the region surrounding the sialic acid binding site in wild-type sialin, whereas in the LO state of R39C sialin, this communication preferentially bypasses this region. Therefore, the R39C mutation may impede the LO to CO conformational transition required for sialic acid transport, providing a plausible mechanistic framework for the decreased transport activity, and possibly the decreased lysosomal localisation, observed in Salla disease.

**Highlights:**

- The R39 residue participates in an interaction triplet, which negatively regulates an ionic lock stabilising the lumen-open conformation
- The R39C mutation is associated with a stronger ionic lock in the simulations, and may favour the lumen-open state
- Correlation network analysis suggests an allosteric coupling between the ionic lock and the region surrounding the sialic acid binding site
- The R39C mutation alters the inferred allosteric coupling between the ionic lock and the region surrounding the sialic acid binding site

**Graphical abstract:** 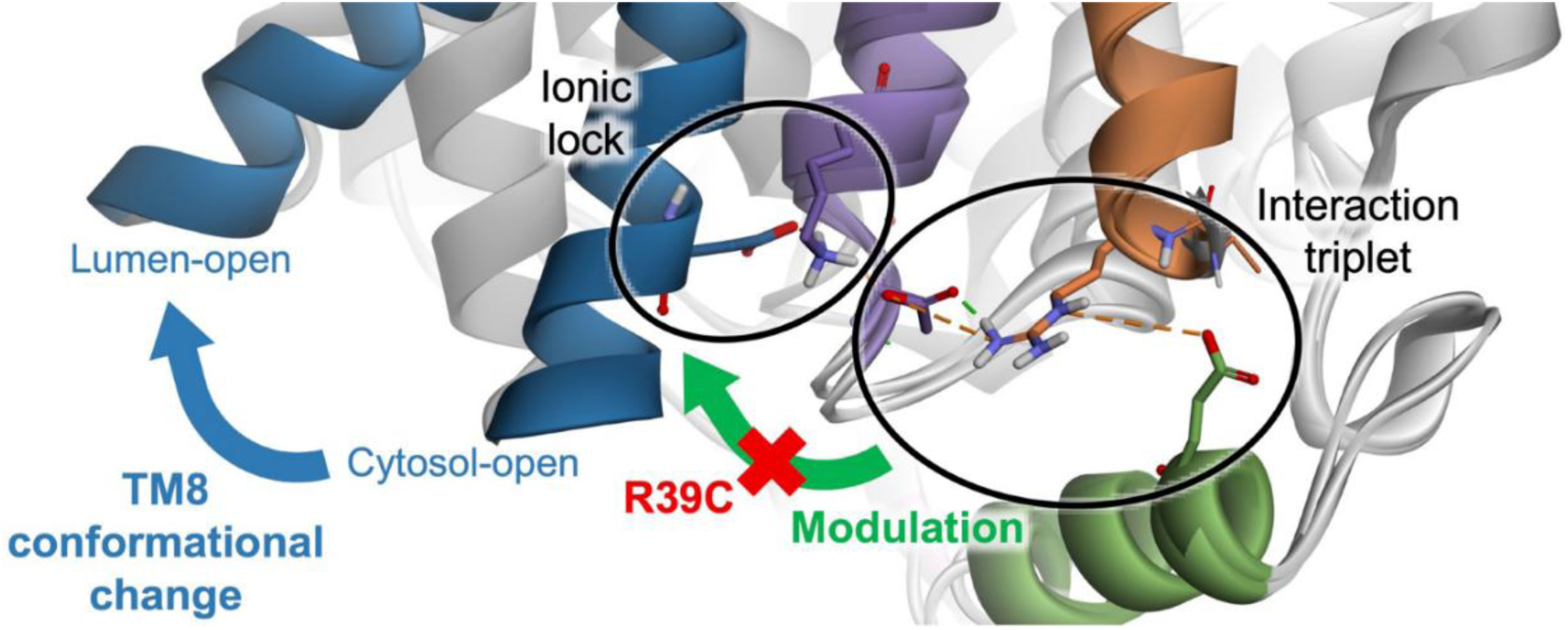

## Introduction

Free Sialic Acid Storage Disorders (FSASD) are rare, autosomal recessive diseases caused by mutations in the *SLC17A5* gene, which encodes sialin, the lysosomal exporter for sialic acids and acidic hexoses [1,2]. The most common, milder form, Salla disease (SD), is predominantly caused by the hypomorphic R39C mutation, which reduces the transport activity of sialin and impairs its delivery to lysosomes [1,3,4]. This leads to abnormal accumulation of sialic acid in lysosomes [5,6], disturbances of ganglioside homeostasis [7–10], hypomyelination [11,12], and severe neurological deficits [13]. Patients typically present with hypotonia, motor delay, intellectual disability, and progressive neurological decline, for which no treatment is currently available [14].

In this context, pharmacological chaperones represent a therapeutic option. These drugs are small molecules that bind and stabilise mutant proteins, thereby rescuing folding, trafficking and functional activity [15]. Using 3D homology models of human sialin and virtual high-throughput screening, we identified two families of sialin ligands with micromolar affinity [16,17]. One of these ligands, LSP12-3129, rescues the trafficking defect of human sialin R39C. However, it failed to reduce sialic acid accumulation in cells from SD patients because this orthosteric ligand also inhibits sialic acid transport. Allosteric ligands offer a second-generation strategy [18], stabilising the mutant protein conformation while avoiding the drawback of intrinsic lasting inhibition. Such ligands can not only rescue trafficking defects but can also activate transport by favouring rate-limiting conformations, as recently shown for the glutamate transporter EAAT2 [19]. As a first step towards the search of allosteric pharmacological chaperones to treat SD, we decided to characterise the precise molecular defects induced by the R39C mutation in human sialin.

Sialin (SLC17A5) is a membrane transporter that belongs to the Major Facilitator Superfamily, characterised by 12 helices (TM1-12) divided into two bundles, the N- (TM1-TM6) and the C-terminal (TM7-TM12) bundles [20,21]. The co-transport of sialic acid and a proton operates by a rocker-switch mechanism, which alternately exposes the substrate-binding site to either the lysosomal lumen or the cytosol [20]. We have previously generated homology models of the lumen-open (LO) and cytosol-open (CO) conformations based on DGoT [22] and VGLUT2 [23] experimental structures. Subsequently, and more recently, cryo-EM structures of both sialin conformations were published [24,25]. These studies suggested that the interaction of R39 with surrounding residues stabilises the correct folding or the cytosol opening of sialin [24,25]. However, the precise effect of the R39C mutation and the reason why it leads to decreased transport activity is not completely understood.

In this study, we generated new models of apo-sialin in the LO and CO conformations using the recently published cryo-EM experimental structures combined with modelling of the missing loops and termini. We used Molecular Dynamics (MD) simulations alongside mutagenesis studies to understand the importance of the allosteric site surrounding R39 and the changes in the local and global conformation of sialin brought on by the pathogenic mutation R39C. Simulations of the LO and CO conformations in wild-type and R39C sialin revealed key structural changes induced by R39C. Through the analysis of the allosteric communication in the protein, we also show how the R39C mutation affects the entire protein and perturbs the propagation of key allosteric changes. This work provides a novel dynamic picture of the molecular defects causing SD within human sialin.

## Results

### The structure of the R39 site

As previously reported [24, 25], sialin is composed of two N- and C- terminal bundles made up of helices TM1 to TM6 and TM7 to TM12, respectively, which undergo alternating-access rotations during sialic acid transport. To understand the impact of the pathogenic mutation R39C causing Salla disease, we modelled sialin in its LO and CO states. We used recently published cryo-EM structures [25] as a starting point to model the LO (PDB ID: 8U3F) and CO (PDB ID: 8U3D) states. The N- and C-termini and the long loop between TM1 and TM2 (D69-N95) (Figure 1A, pink) were not resolved in the cryo-EM structures. We used a hybrid approach employing homology modelling using AlphaFold2-predicted [26] structures of sialin as templates to model these regions. Because of their high flexibility, we generated an ensemble of 1000 models for each state, keeping the experimentally resolved coordinates fixed and sampling different conformations of the termini and TM1-TM2 loop. Due to the low confidence with which AlphaFold2 models the extreme N-terminus, our models start at residue A26. The best model for each state was selected based on its DOPE (Discrete Optimised Protein Energy) score, a statistical potential that evaluates model quality by assessing the likelihood of pairwise atomic distances based on known proteins structures that has been validated for loop modelling [27]. Due to the innate flexibility of the modelled regions, the selected model represents the most probable among the generated conformations, rather than a unique solution, but the use of fixed experimental coordinates for the resolved regions and the subsequent energy minimisation ensures the reliability of the model as a starting point for this study. The model protonation was done, as described in the method section, to approximate the pH gradient between the lysosomal lumen and the cytosol.

**Figure 1.**
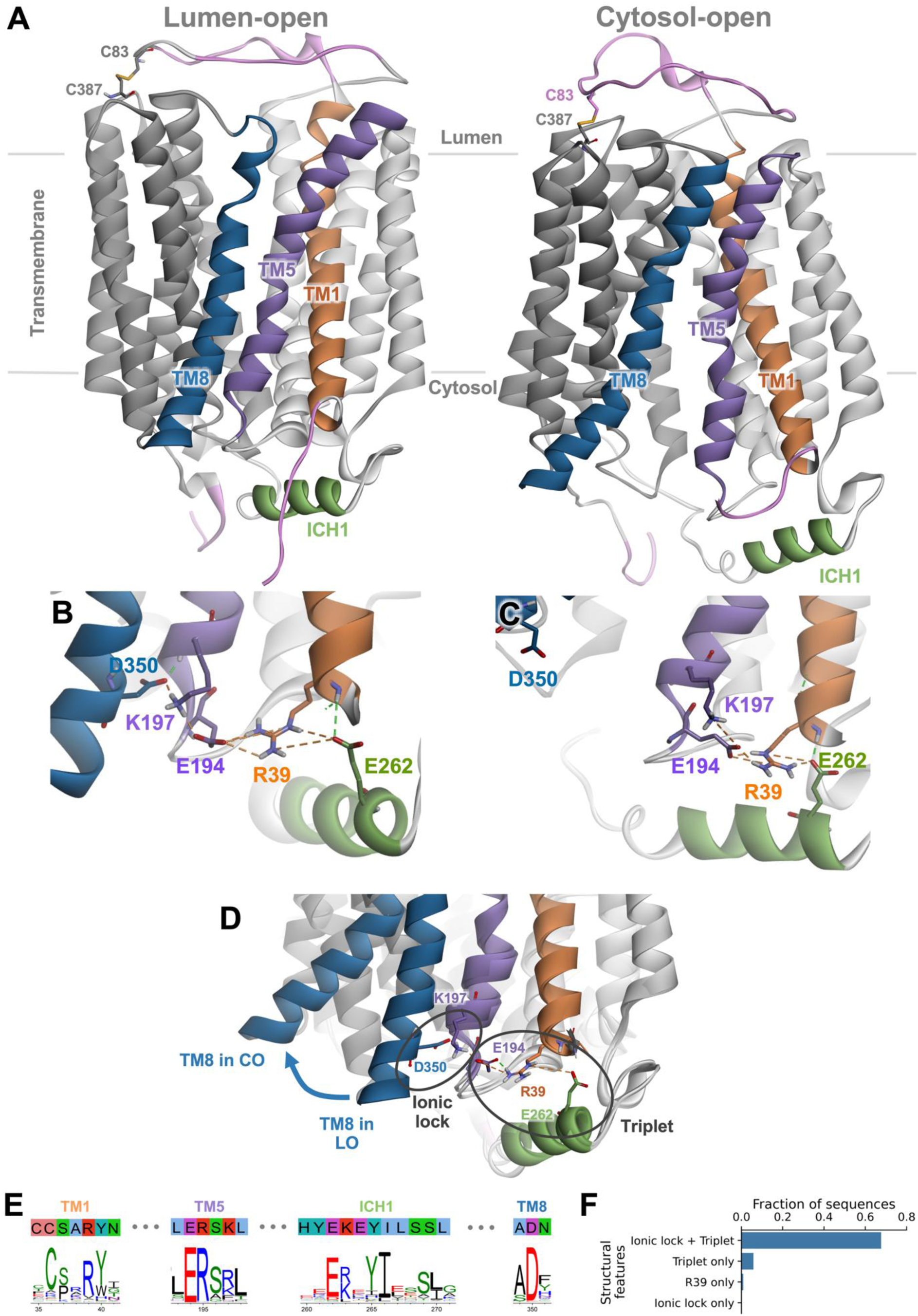
Modelling and analysis of the R39 site of sialin. **A)** The completed and minimised models of sialin in the LO (PDB ID: 8U3F, left) and CO (PDB ID: 8U3D, right) conformations are shown. The missing loops and termini (pink) in the experimental models were modelled using a combination of homology modelling and AlphaFold2. The C- (grey) and N-bundles (white) are shown with helices TM1 (orange), TM5 (purple), and TM8 (blue) highlighted. The two bundles are connected over a long loop containing the ICH1 helix (green) in the cytosolic side. Another long loop between TM1 and TM2 connects the two bundles over a disulphide bond (C83-C387) on the luminal side, forming a “lid” region. **B** and **C)** Close-up views of the R39 site are shown for the LO (B) and CO (C) conformations. R39 is shown to interact via hydrogen bonding (green dashed line) and charge-charge (orange dashed line) interactions with E194 and E262. **D)** Comparison of the TM8 tilt. The superimposed, minimised, and completed models of the LO and CO conformations are shown. Residues R39, E194, and E262 form a triplet and residues K197 and D350 form an ionic lock. TM8 tilts and moves away from TM5 (blue arrow) when sialin changes from the LO to the CO conformation. The residues and their interactions are only shown for the LO conformation. **E)** Sequence logo of segments of TM1, TM5, ICH1, and TM8. The sequence logo generated from the multiple sequence alignment of SLC17 family members is shown alongside the sialin sequence segments. **F)** The conservation of the triplet (R39/E194/E262) and the ionic lock (K197/D350) in the multiple sequence alignment of SLC17 family members is shown. The triplet is considered conserved if in the position equivalent to R39 and E194/E262 in sialin, a basic (K/R) and two acidic (D/E) residues are present, respectively. The ionic lock is considered conserved if in the position equivalent to K197 and D350 in sialin, a basic residue (K/R) and an acidic residue (D/E) are present, respectively.

In our hybrid models, the two bundles are connected through two linkers that extend and contract upon transition between LO and CO conformations. On the luminal side, the long loop between TM1 and TM2 (N-bundle) covers the C-bundle, forming a “lid” over the protein (pink in Figure 1A) connected to the C-bundle by a disulphide bond between residue C83 in the lid and C387 in the loop between TM9 and TM10. A similar disulphide bridge between the long TM1-TM2 loop and the C-bundle has been depicted in VMAT2 [28]. On the cytosolic side, the long loop between TM6 and TM7 (residues V248 to L289) links the two bundles further and contains the intracytosolic helix 1 (ICH1, green in Figure 1A). ICH1 is bound to the N-terminal bundle through interactions with residues in TM1 and TM5. This region is of particular interest due to the presence of the pathological mutation R39C.

#### Identification of a conserved triplet and an ionic lock

On the cytosolic side of the N-bundle, R39 forms multiple interactions with surrounding residues, most notably hydrogen bonding interactions/salt bridges with both E194 (TM5) and E262 (ICH1) in the LO and CO models (Figure 1B and C), forming an interaction triplet. R39 also forms apolar interactions and van-der-Waals contacts in both the LO (Figures S1A and B) and CO (Figures S1C and D) conformations, including with residues A38, Y40, N41, L42, and A43 in TM1; L198 in TM5; and Y265 and I266 in ICH1. The large number of van-der-Waals contacts (Figures S1B and D) leads to a tight packing, resulting in R39 being barely solvent accessible (Figures S2A and D) and likely conformationally locked in position. This constraint is further augmented by hydrogen bonds between Y40 and E194 (LO and CO models, Figures S1A and C) and between Y265 and E194 (CO model, Figure S1C), providing further stability to the conformation of E194 and, by extension, R39.

Close to R39, residues I266 and L270 on ICH1 make apolar interactions with proline residues P191 in the loop between TM4 and TM5 and P252 in the loop connecting TM6 to ICH1 (Figures S1A and C). These interactions further anchor ICH1 to the N-terminal bundle. Moreover, these residues are part of a hydrophobic cluster surrounding the triplet on the N-bundle side (Figures S2B, C, E and F).

The environment of R39 also includes residues K197 (TM5) and D350 (TM8) which form a salt bridge in the LO conformation (Figure 1B), contributing to sealing the substrate translocation pathway from the cytosolic compartment. This salt bridge breaks apart in the CO conformation due to the rocker switch movement of the bundles (Figure 1C). It thus constitutes an ionic lock, much like the one observed in G-protein coupled receptors [29]. In our models, we observe the K197-D350 salt bridge in the LO conformation as expected (Figures 1B and C) and, additionally, we see that K197 forms a salt bridge with E194 in both conformations (Figures 1B and C). This series of salt bridges connects the R39 site to TM8 in the LO conformation (Figure 1D).

These observations prompted us to examine whether the R39/E194/E262 interaction triplet and the K197/D350 ionic lock are conserved in the SLC17 transporter family, using a multiple sequence alignment of 7455 non-redundant sequences of SLC17A1-9 across different species. Indeed, several key residues are conserved (Figure 1E) in particular C36, R39, and Y40 in TM1; E194 and R195 in TM5; E262, Y265, and I266 in ICH1; and D350 in TM8 (Figure 1E). R39 is conserved in 74.5% of sequences. While K197 is less conserved, this position is often occupied by an arginine, which would also form a salt bridge. By defining the interaction triplet as an arginine at the position equivalent to R39 in sialin and acidic residues at the positions equivalent to E194 and E262, and the cytosolic ionic lock as a basic (R/K) and an acidic residue at the positions equivalent to K197 and D350, respectively, we found a concurrent conservation of the triplet and the ionic lock in 67.6% of sequences (Figure 1F). Only 5.8% of sequences have a triplet without ionic lock, while 0.2% sequences have an ionic lock without the triplet. The frequent concurrent conservation of the cytosolic ionic lock and the triplet, the spatial proximity and the described interactions suggest these structural features may be potentially mechanistically important and that a cooperative effect between them may be present.

### Experimental evaluation of the R39 site features

We mutated several residues in the R39 site located on TM1, TM5, or ICH1 to test their importance in sialin function. Most residues were mutated to alanine, including a complete alanine scanning of ICH1 from Y261 to L270. Some key residues were also mutated to other amino acids. We also introduced a proline into ICH1 (E264P) to test the importance of its helical character. These mutations were tested for their effect on the lysosomal localisation and the sialic acid transport activity of human sialin.

The lysosomal localisation of sialin was assessed by comparing the intracellular distribution of transiently expressed, EGFP-tagged sialin with that of a lysosomal marker (LAMP1) in HeLa cells (Figures 2 and S3). The colocalisation was quantified by calculating the Pearson correlation coefficient between the pixel fluorescence intensities of EGFP-sialin and LAMP1 (Figures 3A and B). The presence of an arginine at position 39 (R39) is essential for the lysosomal localisation (mutations R39C, R39D, R39F, R39K and R39M), as well as the presence of acidic residues at positions 194 and 262 (mutations E194A, E194 and E262A), in agreement with the existence of an interaction triplet. Breaking the helicity of ICH1 by mutating E264 into a proline disrupts the lysosomal localisation of sialin, whereas the essential role of residues E262 and I266 supports the importance of this helicity. The localisation was not affected by mutations E264A and E264K, in contrast with the proposed dual interaction of R39 with E262 and E264 reported in a cryo-EM structure (PDB ID: 8DWI) [24] and in agreement with E264 pointing away from the two bundles in our LO and CO models (Figures 3E and F).

**Figure 2.**
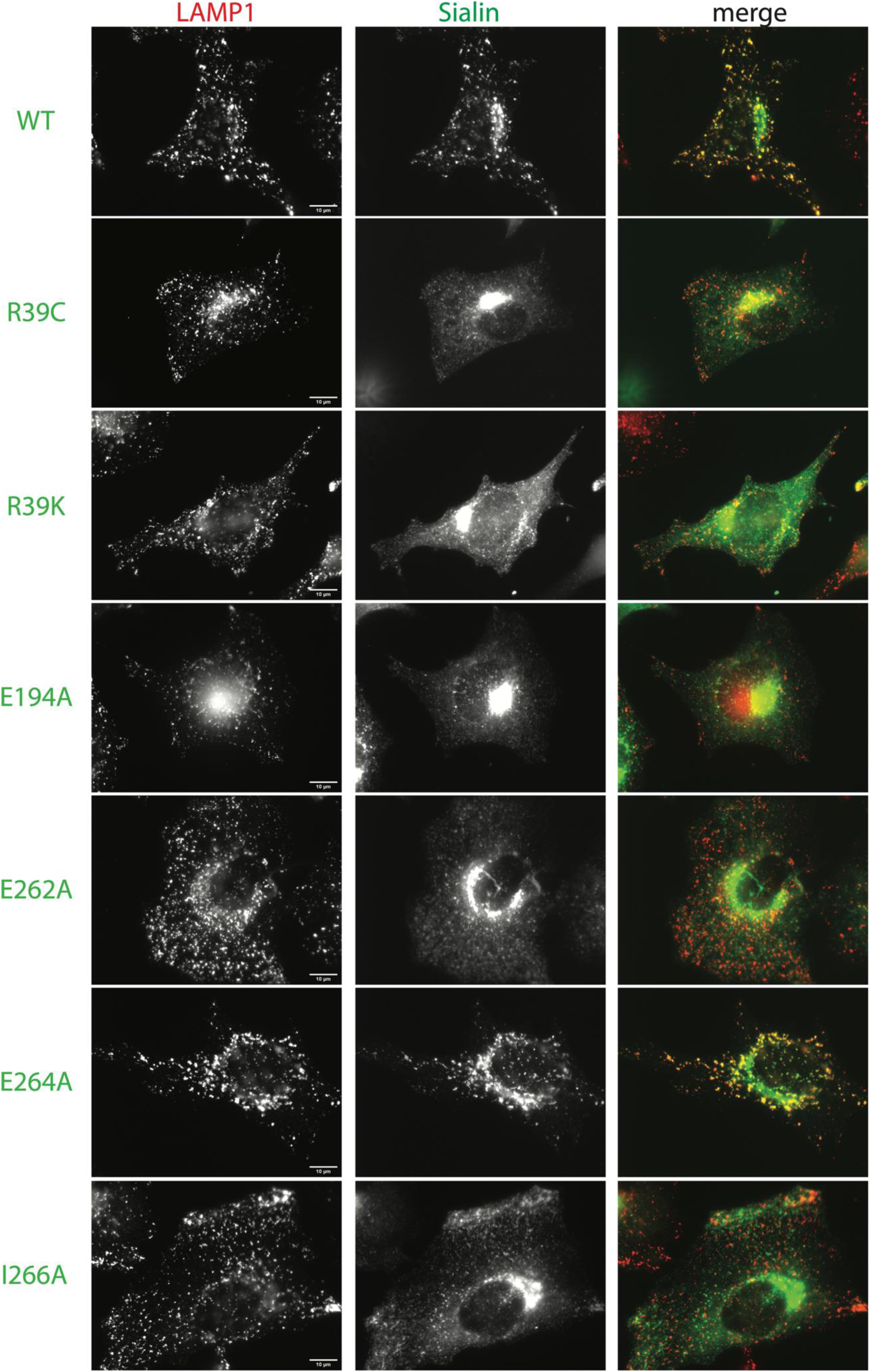
Representative images of the effects of mutations on the lysosomal localisation of sialin visualised by epifluorescence microscopy. Wild-type (WT) or different mutated human sialin (R39C, R39K, E194A, E262A, E264A and I266A) tagged with EGFP (green) constructs were transiently expressed in HeLa cells by electroporation. After two days, cells were fixed and analysed under fluorescence microscopy using LAMP1 immunostaining (red) to detect late endosomes and lysosomes. The scale bar represents 10 µm.

**Figure 3.**
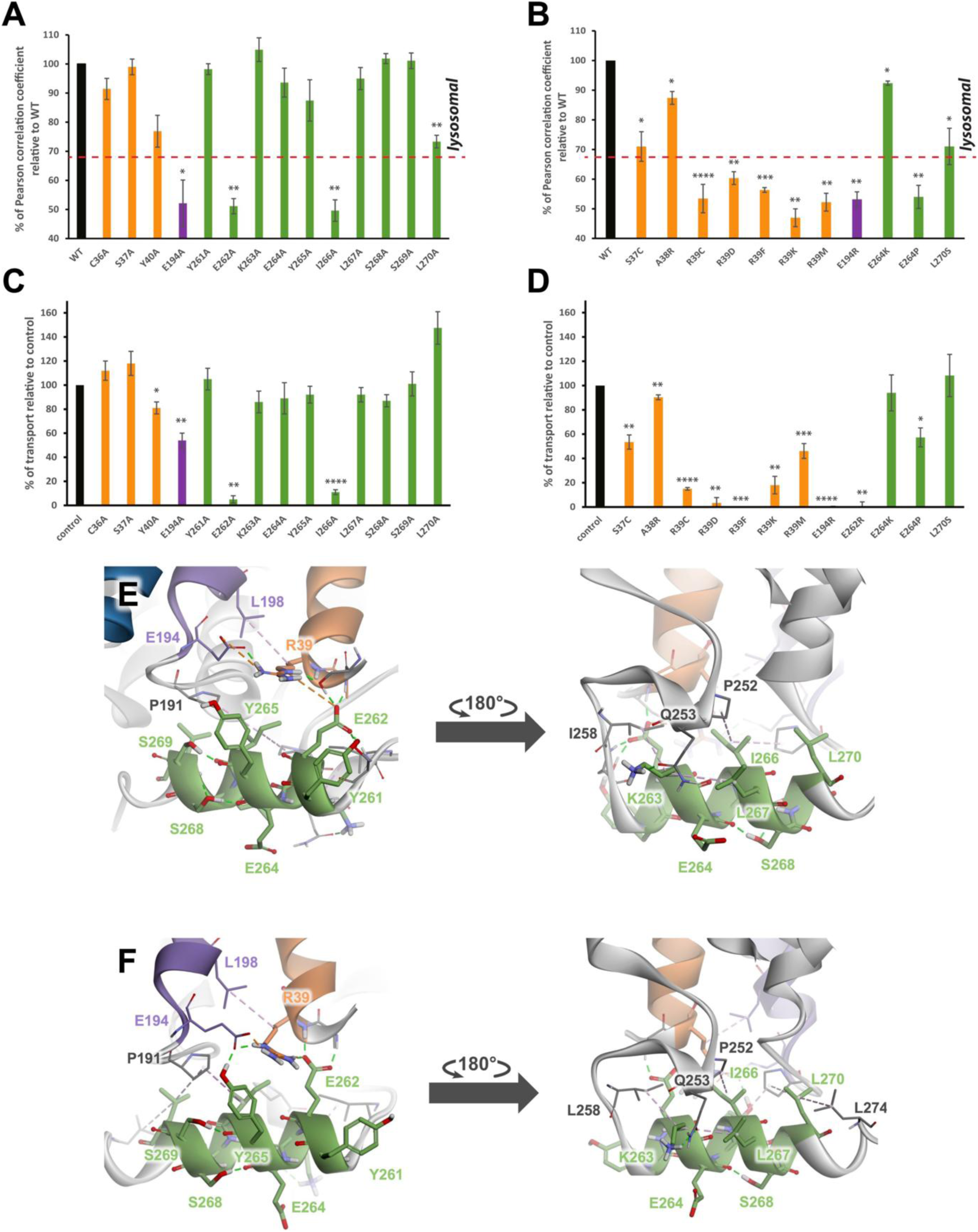
Mutagenesis experiments on the sialin R39 site. **A** and **B)** Wild-type (WT) or different mutated human sialin tagged with EGFP were transiently expressed in HeLa cells by electroporation. After two days, cells were fixed and analysed under fluorescence microscopy using LAMP1 immunostaining to detect late endosomes and lysosomes. In independent experiments, colocalisation was quantified using scatter plots of the sialin and LAMP1 pixel intensities. The graphs show the Pearson correlation coefficients normalised to that of WT (100%) in three different experiments for the mutants, except for R39C, 6 experiments, with 10 cells per experiment. The statistical analysis was performed by a one-sample t-test comparing the mean of the values for each mutant with a hypothetical mean value of 100: *= p ≤ 0.05, ** = p ≤ 0.01, *** = p ≤ 0.001, ****= p ≤ 0.0001). The red dotted line represents a Pearson coefficient of 0.5, above which we consider that sialin is targeted to lysosomes. It is calculated taking into consideration that the mean Pearson correlation coefficient for WT sialin is 0.73. **C** and **D)** Transport activity was measured in a whole-cell assay in which sialin was redirected to the plasma membrane by mutation of the lysosomal sorting motif to facilitate transport measurements (L22G L23G). Human L22G L23G sialin tagged with EGFP with/without mutations was transiently transfected by lipofection in HEK 293T cells. Human sialin (control) and its mutants were assayed for [^3^H]Neu5Ac uptake at pH 5.0 in 3-6 independent experiments. The graphs show the transport activity relative to that of sialin L22G/L23G (100%) The statistical analysis was performed by a one-sample t-test comparing the mean of the values for each mutant with a hypothetical mean value of 100: * = p ≤ 0.05, ** = p ≤ 0.01, *** = p ≤ 0.001, ****= p ≤ 0.0001). **E** and **F)** Close-up views of ICH1 interactions in the completed and minimised models of the LO (E) and CO (F) conformations. Residues in ICH1 (green) are shown to interact with TM1 (orange), TM5 (purple), and other parts (grey) of sialin. Hydrogen bonding (green dashed lines), charge-charge (orange dashed lines), and apolar (purple dashed lines) interactions are shown for residues Y261 to L270 and R39. Hydrogen bonds between backbone atoms involved in the formation of alpha-helices are not shown and only polar hydrogens are displayed to improve visibility. Residues A26-C36 of the N-terminus are not shown to improve visibility.

We also tested the effects of the mutations on the transport activity of sialin, using a sorting mutant (L22G/L23G) redirected to the plasma membrane. In this transport assay, the poorly tractable lysosomal sialic acid export is replaced by a whole-cell import of radiolabelled N-acetylneuraminic acid ([^3^H]Neu5Ac) in acidic medium to mimic the lysosomal lumen [1,16,17]. Prior to the [^3^H]Neu5Ac transport measurements, the surface expression of the EGFP-sialin L22G/L23G constructs in HEK 293 cells was assessed by epifluorescence microscopy (Figures S4, S5 and S6). Remarkably, the effect of the mutations on Neu5Ac transport was highly similar to their effect on lysosomal localisation (Figures 3A to D), suggesting that a common mechanism underpins the two effects. In particular, Neu5Ac transport was abolished by the E262A and I266A mutations and severely reduced by most R39 mutations. Mutation E194R also abolishes Neu5Ac transport and severely impairs lysosomal localisation, in agreement with the interaction triplet. The sole discrepancy between the two assays was the mutation L270A, which significantly decreases lysosomal localisation, but preserves Neu5Ac transport. Of note, L270 selectively interacts with P191 in the CO, but not in the LO, model (Figures S1A and C). This indicates that while most R39 site interactions may be critical for both the trafficking and transport activity of sialin, some of them may play a differential role if they occur in a selective state of the transport cycle.

### Molecular Dynamics simulations show the dynamic character of the R39 site

To go beyond the static image offered by cryo-EM structures, we ran MD simulations on the wild-type sialin in the LO and CO conformations. Each system underwent 91 ns of equilibration, followed by a 450 ns production run performed in triplicate, enabling sufficient sampling of the R39 site conformation. We analysed the last 300 ns of the trajectory, which allowed us to look into distinct conformations of the sites.

#### Non-bonded interactions between R39 and surrounding residues in MD trajectories

The simulations revealed that some interactions remain stable in the wild-type protein. The minimum distances between the heavy atoms of R39 and residues S37 in TM1, E194 in TM5, and E262, Y265, and I266 in ICH1 remain <4.5 Å throughout the trajectories, indicating that these residues are consistently in contact with R39 (Figure S7A). Several R39 interaction patterns were observed in the LO and CO conformations (Figure S8). However, several non-bonded interactions between R39 and E194 and between R39 and E262 remain throughout the simulations (Figure 4A and S9A), in agreement with the drop in lysosomal localisation and transport activity induced by the E194A and E262A mutations (Figure 3). The stability of the R39 residue interactions is also consistent with the mutagenesis data for I266, but not for S37 and Y265.

**Figure 4.**
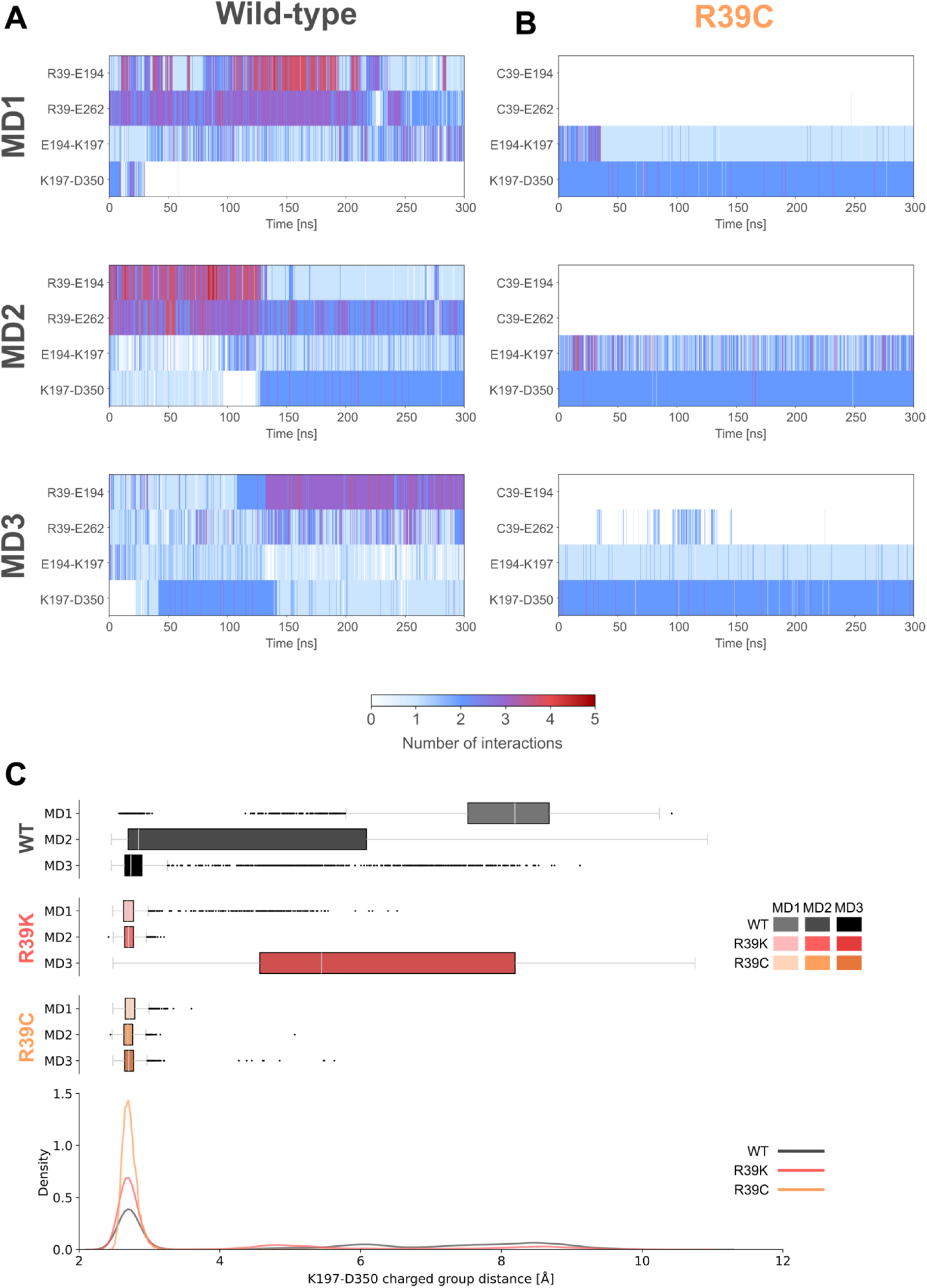
Triplet and ionic lock interaction in wild-type and mutant Sialin. **A** and **B)** Non-bonded interaction monitoring of the wild-type and R39C LO trajectories. The interactions for the C/R39-E194, C/R39-E262, E194-K197, and K197-D350 pairs are shown during MD simulations for wild-type (A) and R39C mutant (B) sialin in the LO conformation. The data of the last 300 ns of the production run of all three replicas, MD1 (top), MD2 (middle), and MD3 (bottom), are shown for each mutant. Interactions are shown as vertical lines, where the colour of the line indicates the number of interactions, ranging from 0 (white) to 5 (red). **C)** Boxplot showing the distance between the functional groups of K197 and D350 (d_4_) in the LO conformation. The minimum distance between the heavy atoms of the amino group of K197 and the carboxylate group of D350 is shown for wild-type (WT, black), R39K (red) and R39C mutant (orange) sialin. The data of the last 300 ns of the production run of all three replicas, MD1 (lighter), MD2 (medium hue), and MD3 (darkest), are shown for each mutant. A density plot showing the K197-D350 distance (d_4_) as a comparison between wild-type, R39K, and R39C mutant is also shown. The data in the density plot includes the last 300 ns of all replicas combined for each mutant.

#### A stabilising hydrophobic cluster

The MD simulations also revealed a dynamic picture of the apolar interactions between residues I266 and L270 and their surrounding proline residues P191 in the TM4-TM5 loop and P252 in the TM6-ICH1 loop (Figures S1A and C). Analysis of the hydrophobic interactions between the pairs I266-P191, I266-P252, L270-P191, and L270-P252 over the three replicas (MD1-MD3) for the LO (Figure S10) and CO (Figure S11) conformation show a constant contact between I266 (ICH1) and the prolines 191 and 252 and mostly no contact between L270 and these prolines. These findings may explain the loss of transport activity in the I266A mutant, but not the L270A mutant (Figure 3C). Additional hydrophobic residues next to P191 and P252, form a hollow pocket around the triplet (Figures S2B, C, E and F) and stabilise the cytosolic environment of R39.

#### Relationship between the triplet and the ionic lock connecting the N- and C-bundles

Looking at residues R39, E194, K197, E262, and D350 during MD simulations revealed much more intricate interactions than the static picture provided by the cryo-EM structures. The distances between the positively charged guanidinium group of R39 and the negatively charged carboxylate group of E194 and E262 fluctuate throughout the trajectories, but remains in most cases within a 2.5-5 Å range (d_1_ and d_2_ in Figures 5, S12, and S13). Such short distance allow the formation of hydrogen bonds and/or salt bridges with these residues in the LO and CO conformations (Figure S8), indicating that this triplet is dynamic yet stable in both conformations. In contrast, the distance between the carboxylate of E194 and the K197 ammonium group (d_3_ in Figure 5) fluctuates over a wider range between strong interaction distances (< 4 Å) and weaker to no electrostatic contact (around 9 Å) during the trajectories (Figures 4A, S9A, S12, S13). This leads to different conformations during the simulations (FigureS8) where the triplet ionic interactions are mostly present (Figure 6, solid orange lines) while the salt bridge between E194 and K197 opens and closes (Figure 6, dashed orange line). The movement of the residues enables E194 to interact with K197 in such a way that K197 can form simultaneous interactions with both E194 and D350 in the LO conformation (Figure 4A and S12; Figure S8B and C), that is when the cytosolic ends of TM5 and TM8 are in close proximity. In this LO conformation, the ionic lock between K197 and D350 is thus intermittent (Figure 6, dashed orange lines; Figures 4A and S12). These features strongly suggest that the E194-K197 interaction may function as a relay between the triplet and the ionic lock (Figure 6, dashed orange lines).

**Figure 5.**
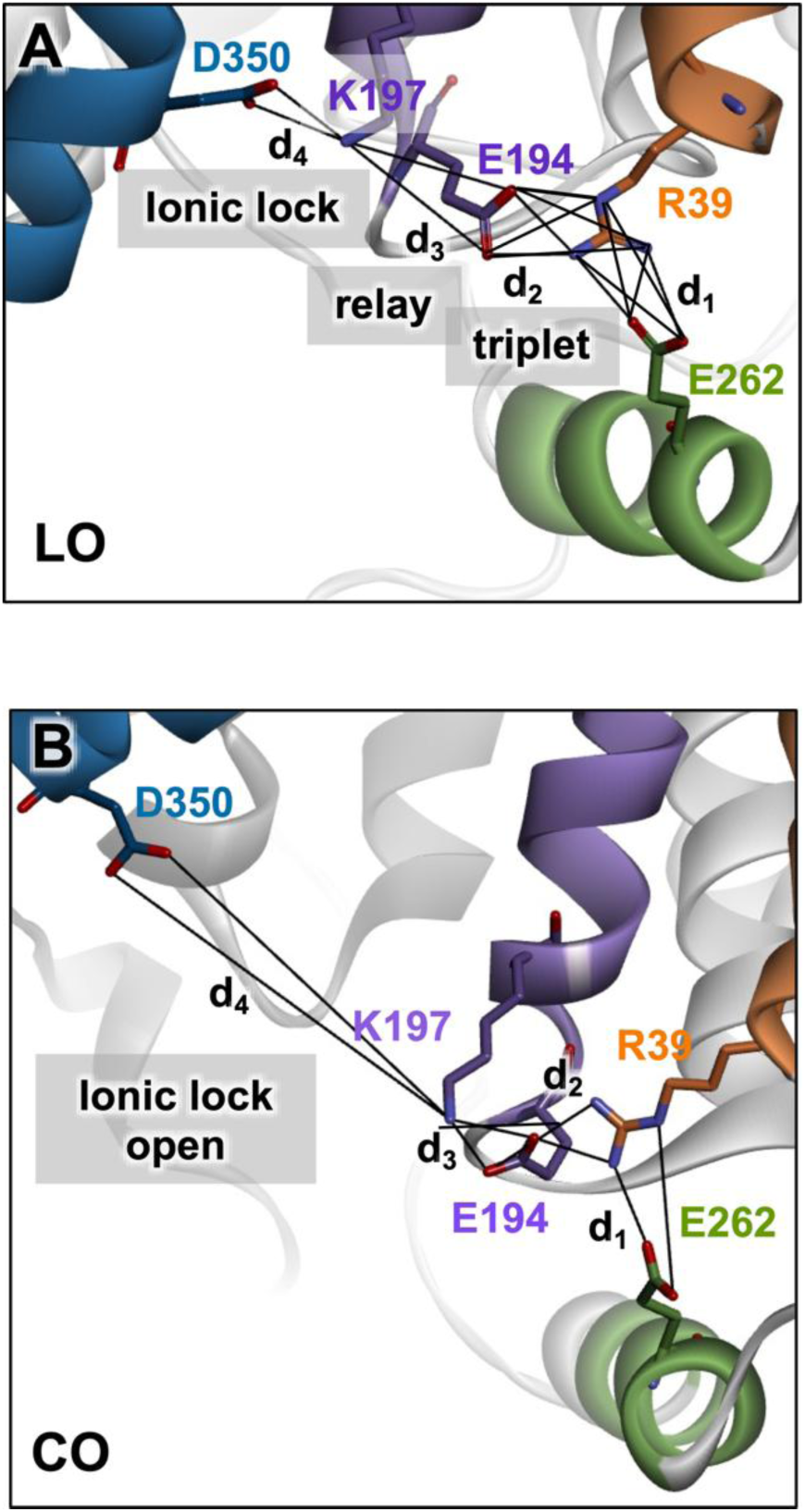
Distances between charged nitrogen and oxygen atoms of the side chains in the triplet, relay and ionic lock in LO **(A)** and CO **(B)** conformations. The distances d_1_, d_2_, d_3_ and d_4_ are the minimal distances between the side chain heteroatoms of E262-R39 (d_1_), E194-R39 (d_2_), E194-K197 (d_3_) and K197-D350 (d_4_). The black lines represent all pairwise distances measured. At each frame, the minimum of these distances was selected to generate Figures S12 and S13.

**Figure 6.**
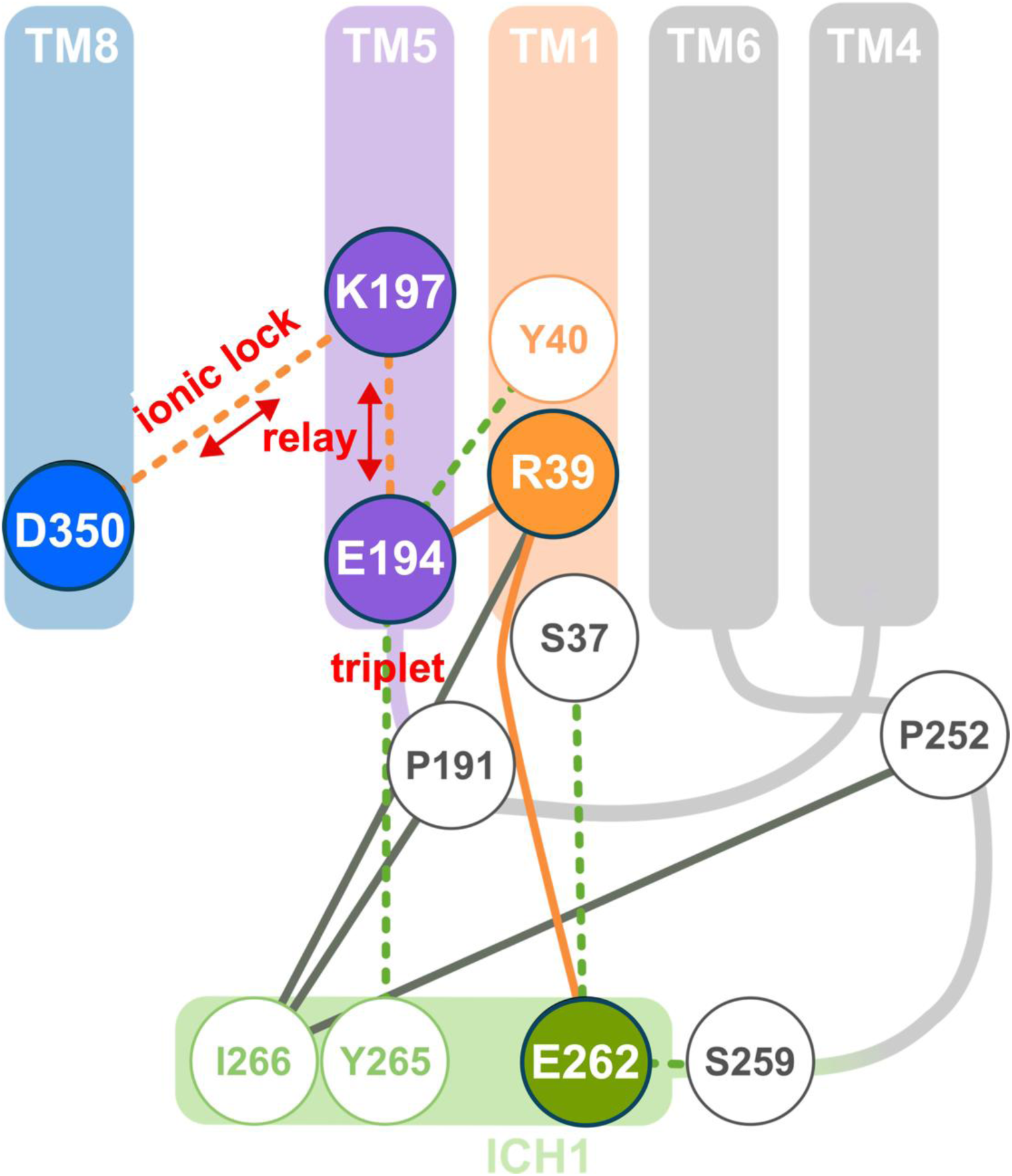
Schematic representation of R39 site interactions in the LO conformation. A schematic representation of the interactions found in the R39 site is shown. The residue R39 on TM1 (orange) interacts through stable attractive electrostatic interactions (orange solid lines) with E194 in TM5 (purple) and E262 in ICH1 (green) making a triplet motif and apolar interactions (grey solid lines) with I266. Fluctuating hydrogen bonding interactions (green dashed lines) between Y40 in TM1 and E194 in TM5, and between E262 in ICH1 and S37 in the N-terminus and S259 in the loop between ICH1 and TM6, stabilise the conformation of the glutamates. Residues K197 in TM5 (purple) and D350 in TM8 (blue) interact, forming the cytosolic ionic lock. E194 interacts with K197, forming fluctuating attractive electrostatic interactions (orange dashed lines) that relay the triplet effect and weaken (red arrows) the ionic lock (K197-D350, interaction, orange dashed line). Residue I266 interacts over stable apolar contacts with P191 and P252.

Monitoring the distances associated with this relay (d_3_), the ionic lock (d_4_) and the triplet (d_1_ and d_2_) during the MD trajectories shows that the ionic lock is disrupted (d_4_ ≥6 Å) when the d_2_ distance is short (d_2_ ∼ 3.0 Å) independently of the relay, provided that the relay maintains an interaction as observed in simulations MD1 and MD2 of WT LO conformation (Figure S12). In this case, the electrostatic influence of the triplet is transmitted to K197 and the ionic lock. On the other hand, the ionic lock is closed when the relay interaction is broken (with long d_3_ ∼7 Å) disrupting the withdrawing effect of R39 as observed in MD3 of the LO conformation (between 150 and 300 ns, Figure S12) and when the relay interaction (d_3_) is strong but the triplet (d_2_) is too weak and the modulating effect is not efficient (MD2 and MD3, Figure S12). Throughout the trajectories E262 remains within a short range (<5 Å) from R39, indicating a stable interaction, which is likely adding the positioning of R39 in a way where it can participate in the above described interactions. These observations suggest a potential mechanism, summarised in the scheme of Figure 6, in which R39 stabilises the position of E194, enabling its interaction with K197 in a manner that weakens the D350-K197 ionic lock. This facilitates its disruption, promoting the transition from the LO to the CO conformation. Therefore, the triplet interactions of R39 appear to modulate the ionic lock to regulate the LO to CO transition during the transport cycle (Figure 6).

The interactions between the triplet and the cytosolic ionic lock are further facilitated by the hydrogen bonding interactions of E194 and E262 with Y40 and Y265 (Figure 6 and S8). These hydrogen bonds are variable across replicas and are found to be present in the different configurations of the ionic lock/relay/triplet interactions, indicating that they might play a secondary stabilising role.

### The Salla disease R39C mutation interrupts key functional interactions in sialin

To study the effects of the Salla disease mutation R39C, we ran simulations of this mutant in both the LO and CO conformations. The same starting coordinates of the protein model, membrane, and solvent molecules as in the wild-type simulations were used for each state, differing only in the mutated residue. The same simulation protocol was also employed, and all simulations were run in triplicate.

In the R39C simulations, the lack of an arginine at position 39 leads to a complete loss of the electrostatic interactions with the glutamate residues E194 and E262 of the triplet in the LO (Figures 4B, 7C) and CO (Figures S9B, S14B) conformations. The cysteine at position 39 cannot form attractive interactions with these glutamates (Figures 7C and S14B), leading to a loss of contacts between ICH1 and the TM1 and TM5 helices of the N-terminal bundle. This is also evident in the distances between R39 and other ICH1 residues (Y261-L270, Figure S7), whose distributions are broader and show higher values in the R39C trajectories compared to the wild type in both conformations. The contacts between R39 and the residues Y265 and I266 are particularly affected since the distance distributions are predominantly <4.5 Å and >4.5 Å in the wild type and R39C proteins, respectively (Figures 7 and S7). ICH1 is thus able to move more freely in the mutant.

**Figure 7.**
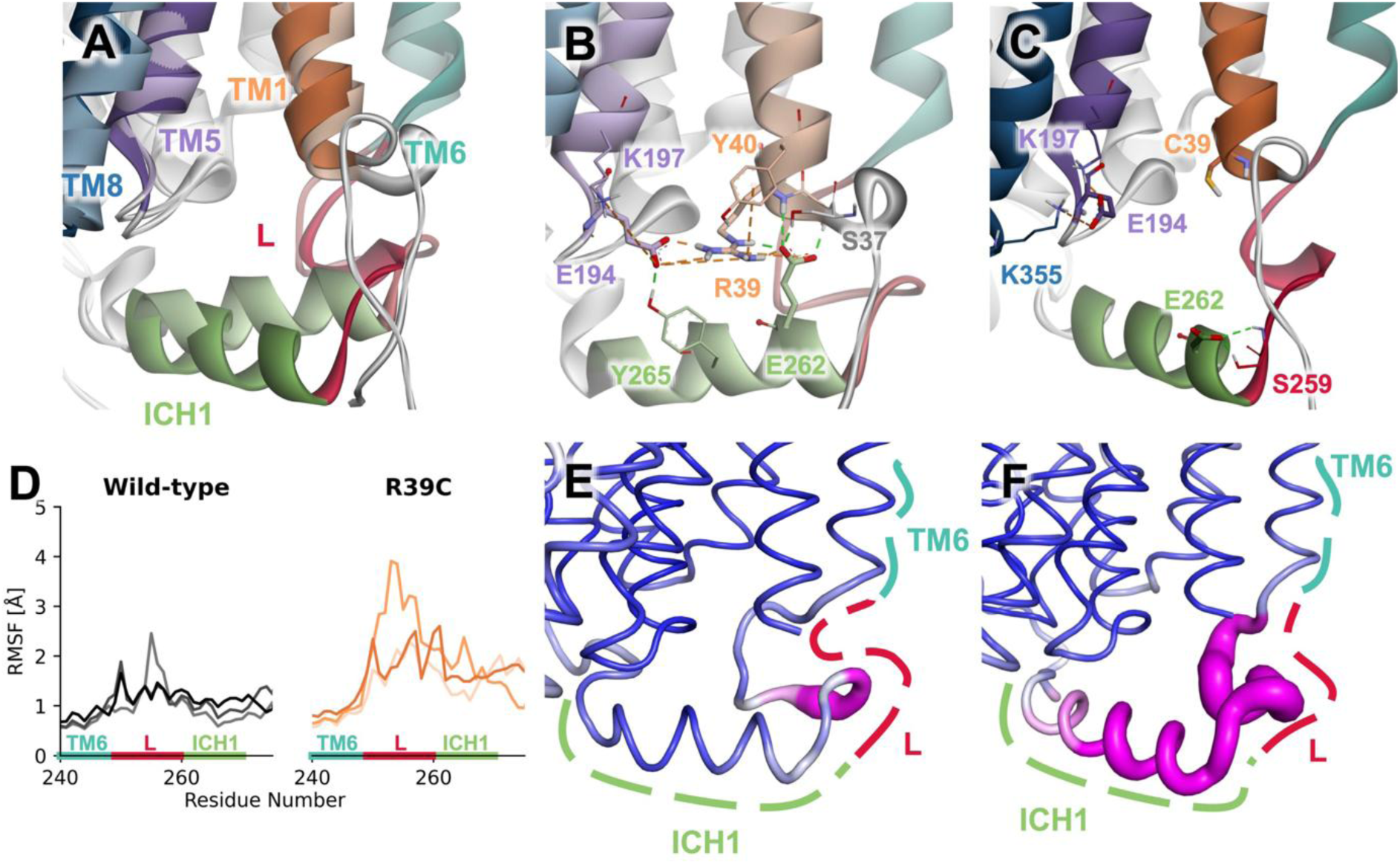
Differences in R39 site interactions between R39C mutant and wild-type sialin in the LO conformation. **A)** Superimposed close-up images of representative structures of wild-type (lighter colours) and R39C mutant (darker colours) in the LO conformation. The movement of ICH1 (green) away from TM1 (orange) and TM5 (purple) is shown in the wild-type and R39C mutant of sialin. **B** and **C)** Interactions of the R39 site in wild-type (B) and R39C mutant (C) shown on representative structures of the LO conformation. In the wild-type, R39 is able to form hydrogen bonds (dashed green lines) and salt bridges (orange dashed lines) with E194 and E262, while in the R39C mutant, these interactions are not possible. **D)** Line plot showing the RMSF values of the Cα atoms of the end of TM6 (teal), TM6-ICH1 loop (L, red), and ICH1 (green) in the wild-type (left) and R39C mutant (right) sialin during MD simulations of the LO conformation. The data shown are from the three replicas: MD1 (lighter), MD2 (medium hue), and MD3 (darkest). **E** and **F)** The RMSF of the Cα atoms is shown mapped on the average structures of wild-type (E, MD1) and R39C mutant (F, MD2) sialin LO trajectories. High RMSF values are shown as thick and magenta, and small ones as thin and blue.

The stabilising hydrogen bonding interactions between the glutamate residues E194 and E262 and surrounding tyrosine and serine residues, respectively, are also affected by the R39C mutation. In the case of E194, the percentage of frames where it interacts with neither Y40 nor Y265 consistently increases in both LO and CO trajectories (Figure S15A). Residue E262, on the other hand, shows a preference for a hydrogen-bonding partner to S259 over S37 in the R39C mutant (Figures 7C, S14B and S15B). The apolar interactions between I266 and the residues P191 and P252 are also less frequent in the R39C mutant than in the wild type in both conformations (Figures S10 and S11).

The hydrophobic contacts, charge-charge, and hydrogen-bonding interactions provide a stable anchoring of ICH1 to TM1 in wild-type sialin, which is reflected in the distances between the Cα atoms of R39 and E262. These distances remain stable in a range from 8 to 11 Å across most replicas in both the LO (Figure S16A) and CO (Figure S16B) states, except for the MD1 CO simulation which shows a stable distance but at higher values (11-13 Å). In the R39C mutant, all replicas show high fluctuations with these distances taking values between 7 and 14 Å in the LO (Figure S16A) and CO (Figure S16B) conformations. Beyond this specific interaction, the overall mobility of ICH1 (residues 261-270) and the adjacent loop connecting TM6 to ICH1 (L) also increases in the R39C mutant, as demonstrated by the Cα atom root mean square fluctuation (RMSF) values of ICH1 and L in the LO (Figures 7D to F and S17A) and CO (Figures S14C and D, S17B) conformations. Altogether, the most affected region is clearly the TM6-ICH1 loop (L in Figures 7 and S14) in the R39C mutant (Figures S17A and B).

Another striking effect of the R39C mutation is the strengthening of the cytosolic ionic lock in the LO conformation (Figure 4A and B). The lack of an intact R39/E194/E262 triplet causes an increased stability of the K197-D350 salt bridge in that conformation. In agreement with the interplay between the triplet and the ionic lock in the wild-type protein, this effect may arise from this lack of interactions between residue 39 and E194, which would otherwise maintain E194 in a conformation capable of weakening the K197-D350 interaction. This observation is consistent with the WT behaviour, where the ionic lock is closed when the triplet interactions are too weak or absent regardless of the strength of the relay (Figure S12). Whereas the distance between the ammonium of K197 and the carboxylate of D350 is quite variable in the wild type, ranging from 2.5 Å to >10 Å (Figures 4C and S12), this distance falls into a tight range of 2.5 to 3 Å in the R39C mutant (Figure 4C), indicating a very strong salt bridge confirmed in the monitoring of non-bonded interactions (Figure 4A and B).

Taken together, our MD simulations show that the R39C disrupts the triplet interactions, resulting in a strengthening of the ionic lock in the LO conformation and, in turn, a stabilisation of this LO conformation. This effect may impede the transition from the LO to the CO conformation in the mutant, resulting in a slower transport activity.

### The artificial R39K mutation does not restore the triplet/ionic lock interaction

To test further this scenario, we studied the effect of the conservative R39K mutation, in which the R39 guanidinium group is replaced by an ammonium group. This mutation impairs both the lysosomal localisation and the transport activity of sialin to an extent similar to that of the pathogenic R39C mutation (Figures 3B and D). This indicates that a positive charge alone on residue 39 is not enough to produce the interactions necessary for proper transporter function and lysosomal localisation. We thus ran simulations on the R39K mutant in both conformations to understand why. As in the R39C simulations, the initial coordinates of the system where the same as in the wild type with only the mutated residue differing.

Comparison of the triplet distances in the wild type (d_2_ R39-E194 and d_1_ R39-E262) and R39K mutant (d_2_ K39-E194 and d_1_ K39-E262) shows that strong simultaneous interactions between K39 and the two glutamates can still occur in the R39K mutant (Figures S18E and H and S19E and H). However, these triplet distances between the charged groups of K39 and E194 and E262 span a broader range of 2.5-10 Å in the mutant, leading to much weaker triplet interactions compared to the wild type where distances remain within 2.5-6 Å. This difference is even more evident when comparing the distributions (edge plots in FigureS18) of the distances K/R39-E194 and K/R39-E262. In the R39K mutant, the density at shorter distances (<4 Å) decreases relative to the wild type. In the LO state particularly, conformations where K39 interacts with either E194 (Figure S18F) or E262 (Figure S18G), but not both simultaneously, become more prevalent in this mutant. This is reflected by the higher density observed in the contour plot (left in Figure S18 and Figure S19 for CO state) at areas where one of the two distances increases (>6 Å) but not the other. Furthermore, in the wild type, E194 stays within a range of 2.5-6 Å from R39 during the trajectories, even while interacting with K197 (Figures S18 and S19). This is not the case in the R39K mutant, where conformations where E194 interacts with K197, but not K39 simultaneously, occur more frequently. This is indicated by the higher densities present in the contour plot (Figures S18 and S19) at areas where the E194-K197 distance is <4 Å and the R/K39-E194 distance is increased (>6 Å). This is especially evident in the LO conformation (Figure S18G). These observations are consistent with the analysis of non-bonded interactions, which show fewer simultaneous interactions between K39 and E194 or E262 (Figures S20 and S21) in the R39K mutant compared to the wild type. In this mutant, the ionic lock of the LO conformation remains closed most of the time in the MD1 and MD2 simulations. In MD3, the ionic lock opens in the beginning of the trajectory (Figure S20B), yet this opening is less frequent than in the wild type (Figure S20A).

Taken together, these simulations show that although the R39K mutation partially restores the triplet interactions as compared with the R39C mutant, the electrostatic interaction of K39 with E194 is not strong enough to pull K197 away from D350 and weaken the K197-D350 cytosolic ionic lock in the LO conformation. The ionic lock is thus strengthened by the R39K mutation as compared with the wild type, impairing the transition from the LO to the CO conformation and slowing down the transport activity similarly to the R39C mutant.

### The R39C mutation affects allosteric communication in sialin

To understand the global effects of the R39C mutation on sialin, we calculated dynamic cross-correlation (DCC) matrices for the wild type and R39C mutant in the LO and CO conformations. The matrices were created by calculating the correlation of the movement of all pairs of Cα atoms for each replica (MD1-MD3) in the last 300 ns of the production run. The correlations range from +1 (perfect correlation), to 0 (no correlation/no coordination), to -1 (perfect anticorrelation) and show how coordinated the movement of two residues is. High coordination is indicative of allosteric coupling between the residues. A consensus matrix (Figure 8) was calculated for each condition (mutant and conformation) by retaining only correlations that are |r|>0.3 in all three replicas and setting the rest to 0. The three resulting filtered DCC matrices with the retained correlations were averaged to yield the consensus matrix (see supplementary files). A threshold of 0.3 was chosen since it allowed for >99% network connectivity across all conditions in the respective correlation networks.

**Figure 8.**
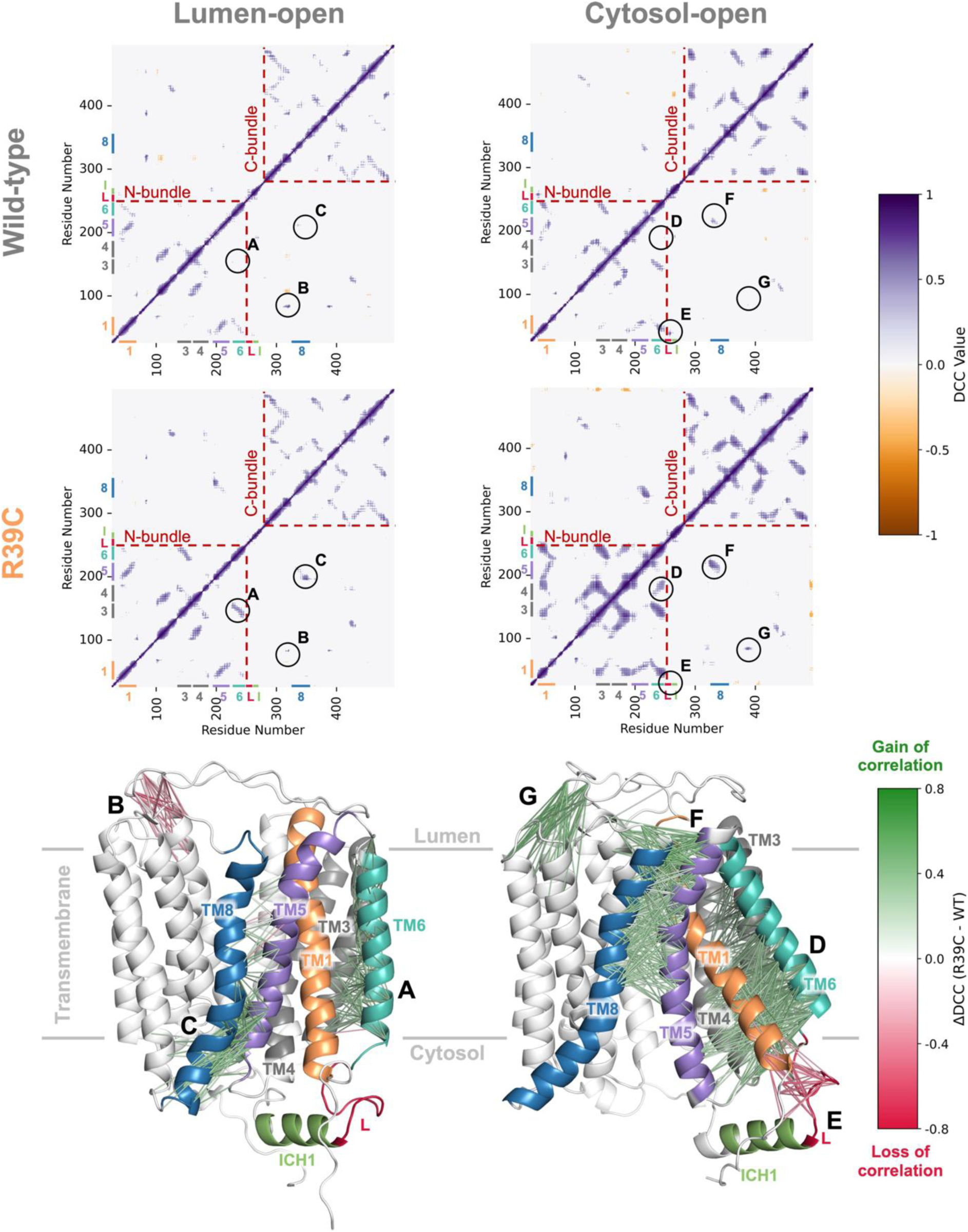
Changes in allosteric communication between wild-type and R39C mutant sialin. Heatmaps showing the consensus dynamic cross-correlation matrices (DCCM) for the LO (left column) and CO (right column) conformations of wild-type and R39C mutant of sialin. Colour scale represents correlation coefficients from -1 (perfect anti-correlation, orange) to +1 (perfect correlation, purple). TM1 (1, orange), TM3, (3, grey), TM4 (4, grey), TM5 (5, purple), TM6 (6, teal), the ICH1-TM6 connecting loop (L, red), ICH1 (I, green), and TM8 (blue) are shown on the x- and y-axes of the DCCM heatmaps. Intra-bundle correlations are denoted as areas within the box of red dashed lines. Structural mapping: Selected differences in DCCM regions (A–G) are visualised on LO (bottom left) and CO (bottom right) structures, with lines connecting Cα atoms representing loss (red) or gain (green) of correlation in the R39C mutant compared to wild-type sialin. LO Conformation: (A) TM3 (grey)–TM6 (teal) correlation increase; (B) Lid–TM7 luminal end correlation decrease; (C) TM5 (purple)–TM8 (blue) cytosolic end correlation increase. CO Conformation: (D) TM4 (grey)–TM6 increase; (E) TM1 (orange) cytosolic end– TM6-ICH1 loop (L, red) decrease; (F) TM5–TM8 luminal end increase; (G) Lid–TM9-TM10 loop increase. The consensus DCC matrices were generated by averaging the matrices from each of the three replicas (MD1-MD3) for each condition after setting correlations that were |r|<0.3 in at least one replica to zero.

#### Correlated movements altered by the R39C mutation

The number of correlations present in the R39C mutant is higher than in the wild type for both the LO (Figure S22A) and CO (Figure S22B) conformations. This is particularly evident in the CO conformation where there are notably more intra-bundle correlations (red dashed lines in Figure 8) in the mutant compared to wild type. This increase indicates a more diffuse allosteric coupling within the bundles and, by extension, a more connected network of allosteric communication, which may allow for distant inter-residue communication to go through alternative, shorter, and potentially non-native, pathways that may not be functional. Considering the lower transport activity of R39C, this increased connectivity may be detrimental to the correct functioning of the transporter.

Selected regions (A-G in Figure 8) of positive correlations in the DCC matrices that differ in the mutant and wild type were further studied to understand how the R39C mutation affects allosteric communication. In the LO conformation, the correlations between TM6 and TM3 (A in Figure 8) increase in the R39C mutant compared to the wild type, likely reflecting the changes in ICH1 and the TM6-ICH1 connecting loop (L) which propagate to the adjacent TM6. The correlation of the movement of the cytosolic ends of TM5 and TM8 (C in Figure 8) also increases in the R39C mutant, which is due to the strengthening of the K197-D350 ionic lock between these helices. The global structural consequence of this strengthening is the increased opening of the aqueous cavity to the lysosomal lumen, with average angles between the main axes of the N- and C-terminal bundles of 20.84±2.42° and 24.39±2.43° in wild-type and R39C sialin, respectively (Figure S23A and B). There are also decreased correlations between the lid region and the luminal end of TM7 (B in Figure 8) in the mutant compared to the wild type, which further highlights the global changes caused by the R39C mutation.

In the CO conformation, correlations are increased between TM6 and TM4 (D in Figure 8), instead of TM6 and TM3 in the LO conformation. Such strengthened dynamic coupling between TM6 and another helix of the N-terminal bundle may again reflect the changes in the adjacent L and ICH1 structural elements. The luminal ends of TM5 and TM8 (F in Figure 8) are also more correlated in the mutant in the CO conformation, similarly to the cytosolic ends of these helices in the LO conformation. The lid region and the TM9-TM10 loop (G in Figure 8) also show an increased correlation in the mutant, highlighting the global perturbations induced by R39C. Decreased correlations are also found between the TM6-ICH1 loop (L) and the cytosolic end of TM1 (E in Figure 8), which are most likely due the increased flexibility of this loop in the R39C mutant as shown by the increased RMSF values (Figure S17B).

#### Network analysis of the correlated movements

The positive correlations of the consensus matrix were used to construct a correlation network. This analysis represents the protein as a correlation network in which residues are connected through correlated motions (Figure S24) [30]. By mapping how movements in one protein region correlate to movements in another, we can trace paths through which allosteric communication passes. The shortest paths between all pairs of residues, corresponding to the highest correlations, provide a way of finding hubs of allosteric communication common to several paths by calculating the betweenness centrality of the residues, that is how often they contribute to the shortest path between two other residues. A high betweenness centrality indicates, therefore, that a residue acts as a bridge/relay along routes of allosteric communication (Figure S24).

We calculated the betweenness centrality of each generated network, normalised it to values between 0 and 1 to make it comparable across mutants and conformations, and mapped it to a corresponding structure of sialin to identify key regions of allosteric communication and how they are affected by the R39C mutation.

In the wild-type LO conformation, the normalised betweenness centrality (NBC) is high in residues lining the central aqueous cavity close to the sialic acid binding site (Figure 9A). These residues are located in both the N-terminal bundle (F47 and Y54 in TM1 and L208 and V211 in TM5) and the C-terminal bundle (C341, Y335, and W339 in TM8 and T397 in TM10). They form a communication hub between the two bundles that is likely central to propagate allosteric effects from one bundle to the other. This is particularly evident when calculating the 500 optimal and suboptimal paths between the residues K197 and D350 that form the cytosolic ionic lock (Figures 9B and I). These paths show a high convergence, always following a similar route along the cytosolic end of TM5 passing, in most cases, through V211 and W339, and the cytosolic end of TM8, with an average path length of 10.58 residues. These paths couple the ionic lock to the sialic acid binding site.

**Figure 9.**
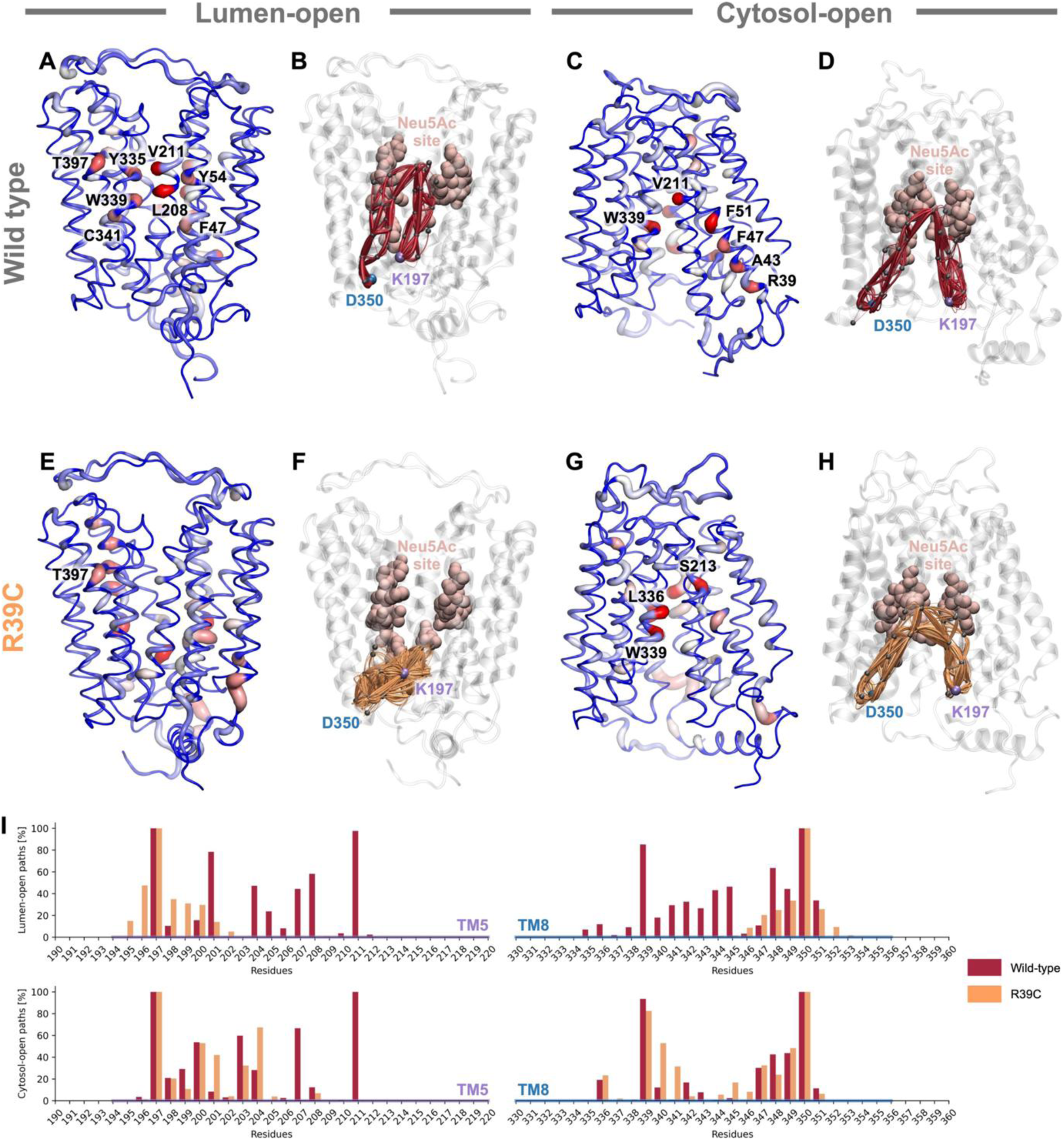
Correlation network analysis in wild-type and R39C. **A**, **C**, **E**, and **G)** Normalised betweenness centrality mapped onto sialin structures in the LO (A and E) and the CO (C and G) conformations for wild-type (A and C) and R39C (E and G) mutant. Thickness and colour indicate normalised betweenness centrality values (red/thick=high; blue/thin=low). **B**, **D**, **F**, and **H**) Suboptimal and optimal paths between residues K197 and D350 involved in the cytosolic ionic lock. The 500 optimal and suboptimal paths between K197 (purple sphere) and D350 (blue sphere) are shown for the wild-type (B and D, red) and R39C (F and H, orange) mutant in the LO (B and F) and CO (D and H) conformations. In the wild-type, the paths (red lines) pass close to the sialic acid binding site (pink) and show high convergence, passing through the same nodes (smaller grey spheres). The R39C mutation alters the allosteric communication between K197 and D350, resulting in shorter paths (orange lines) in both conformations, particularly in the LO conformation. **I)** Barplot showing the frequency of residue participation in optimal and suboptimal paths between K197 and D350. For each residue, the percentage indicates its presence across 500 calculated paths between K197 and D350 in the LO (top) and CO (bottom) conformations in the wild-type (red) and R39C mutant (orange) sialin. Residues belonging to TM5 and TM8 are marked as blue and purple, respectively, along the x axis. The normalised betweenness centrality and optimal and suboptimal paths were calculated using the r>0 values of the consensus DCC matrix. The matrix was generated by averaging the matrices from each of the three replicas (MD1-MD3) for each condition after setting correlations |r|<0.3 in at least one replica to zero. The DCC matrices were generated using the movement of the Cα atoms from the last 300 ns of the trajectories.

In the wild-type CO conformation, the mapping of NBC (Figure 9C) shows a different allosteric communication network. Like in the LO conformation, residues V211 and W339 have increased NBC forming a hub of communication to propagate allosteric changes from one bundle to the other close to the sialic acid binding site. However, residues R39, A43, F47, and F51 in TM1 also show increased NBC in the CO conformation, indicating an increased participation of this helix in the allosteric network and highlighting the importance of R39 in this network. The 500 optimal and suboptimal paths between the residues of the cytosolic ionic lock in the CO conformation follow a similar route as in the LO conformation, following the cytosolic end of TM5, passing through V211 and W339 close to the sialic acid binding site, and going along the cytosolic end of TM8. The average path length is 8.72 residues.

These findings suggest that in both conformations, the breaking and reforming of the ionic lock involve allosteric coupling to the sialic acid binding site, or regions adjacent to it, highlighting the functional connection between the two regions. Conformational changes in the sialic acid binding site (e.g. through the binding of sialic acid) could modulate this allosteric coupling.

The network of allosteric communication in sialin is perturbed by the R39C mutation. In the R39C CO conformation, residue V211 has a decreased NBC value, and residues S213, L336 and W339 have increased values (Figure 9G) relative to the wild type. The increases in NBC of S213 and L336 are likely due to the increased correlations between the luminal ends of TM5 and TM8 (F in Figure 8) in the mutant, which lead to an alternate communication hub between the two bundles away from V211 and towards the luminal side of the transporter. Additionally, TM1 has lower NBC values in the mutant compared to the wild type, indicating a reduced participation of this helix in the allosteric network. These changes affect the optimal and suboptimal paths between K197 and D350 (Figures 9H and I), which are on average shorter (7.83 residues) than in the wild type, with V211 no longer involved in these paths.

In the R39C LO conformation (Figure 9E), the hub residues identified in the wild-type LO conformation show reduced NBC values. Strikingly, the 500 optimal and suboptimal paths between the residues of the ionic lock (Figures 9F and I) are much shorter in the mutant, with an average path length of 5.05 residues passing through the surrounding residues of K197 and D350 without going through the vicinity of the sialic acid binding site. This change is due to the increased correlations (C in Figure 8) between the cytosolic ends of TM5 and TM8 in the R39C mutant, caused by the strengthening of the ionic lock. Consequently, the breaking of the cytosolic ionic lock is disconnected from the sialic acid binding site and its surroundings, potentially contributing to the reduced transport activity of R39C sialin.

Taken together, these findings show that the R39C mutation alters allosteric communication between the two bundles of sialin in both LO and CO conformations, with an uncoupling of the ionic lock from the sialic acid binding site in the LO conformation.

## Discussion

In this study, we combined computational and mutagenesis approaches to investigate the molecular basis of Salla disease (SD). We first generated three-dimensional models of the transporter in the LO and CO states. Our modelling strategy leveraged available cryo-EM structures together with deep learning methods to produce 3D-models of wild-type and mutated sialin that include the regions poorly resolved by cryo-EM. However, we omitted the first 25 residues of the sequence in our models because this extreme N-terminus, which includes the sorting motif responsible for the lysosomal localisation [1,3], could not be modelled with a high degree of certainty.

Our models were protonated in a way that approximates the pH gradient found along the lysosomal membrane. The protonation states of residues were chosen based on p*K*_a_ predictions sampled from MD trajectories to allow for a more dynamic view of the p*K*_a_ variation that occurs as the electrostatic environment of residues changes during the trajectory. While conventional MD simulations maintain fixed protonation states, the approach and computational data presented here provide a well-grounded approximation supported by experimental evidence, acknowledging that model selection and changes in protonation state may influence the electrostatic environment of the protein.

In both our LO and CO models, we found a similar binding motif for R39, which maintains the ICH1 helix connecting the two bundles (TM6-TM7) and the cytosolic ends of TM1 and TM5 in close proximity through salt bridges/hydrogen bonds with E194 and E262. Together, these residues form a triplet motif that is conserved among other members of the SLC17 family including the vesicular glutamate transporter VGLUT (SLC17A6-8) and the sodium/phosphate cotransporter (SLC17A1-4) [22]. Interestingly, in the vesicular nucleotide transporter VNUT (SLC17A9) a related triplet (E90/K91/D210) is also present, but instead connects TM3 to ICH1 as observed in the AlphaFold model [31] (UniProt ID: Q9BYT1). These observations indicate that this triplet constitutes a key structural feature of the SLC17 family and its disruption is, therefore, expected to impair transport activity.

We also identified that a neighbouring ionic lock between residues K197 and D350 in the cytosolic ends of TM5 and TM8, respectively, which is also conserved across members of the SLC17 family. This lock has been already described in the cryo-EM structures [25]. It forms at the cytosolic interface of the bundles in the LO conformation, tightening the two bundles and stabilising the conformation. Such an interdomain ionic lock is present in other transporters of the MFS superfamily, such as the lipid transporter MFSD2A [32,33], the sugar transporter MelB [34–36], and the multidrug resistance transporter MdfA [37–39]. In sialin, upon transition to the CO conformation, these regions of TM5 and TM8 separate while the luminal ends of the bundles get closer, disrupting the lock.

The co-conservation of the triplet and ionic lock motifs and their structural proximity pointed to a potential interaction between these motifs. Indeed, our MD simulations of wild-type, R39K, and R39C sialin revealed a dynamic interplay between these two motifs, which had not been documented by the cryo-EM studies. We observed that the salt bridge between E194 (a component of the triplet) and K197 (a component of the ionic lock) may act as a relay between the triplet and the ionic lock. Residue R39, positioned by its interactions with E262, can interact with E194, which in turn can interact with K197. The E194-K197 interaction thus may transmit the electrostatic effect of the triplet and pulls K197 away from D350, thereby contributing to the opening of the cytosolic ionic lock. The MD simulations suggest that the delocalised positive charge of a guanidinium group can be essential to preserve the modulatory role of the triplet on the ionic lock through the E194-K197 interaction. Accordingly, the replacement of the guanidinium group by an ammonium (R39K mutation) strongly impairs transport activity. When the triplet is disrupted (R39C) or weakened (R39K) by mutations, its weakening effect on the cytosolic ionic lock is lost, leading to strengthening of the ionic lock. This in turn may stabilise sialin in the LO conformation and perturbs the progression of the transport cycle: R39C sialin cannot easily switch from the LO conformation (where it captures sialic acid) to the CO conformation, required to release sialic acid to the cytosol. This interpretation is consistent with a previous study [1] showing that the R39C mutation reduces the velocity (Vmax) of sialic acid transport without altering the affinity of sialin for its substrate.

To further investigate how the R39C mutant affects sialin, we computed dynamics cross-correlated matrices and analysed the resulting correlation networks throughout the protein. In wild-type sialin, the allosteric communication between the N- and C-terminal bundles passes through two key relay points, V211 and W339, in both the LO and CO conformations. These residues V211 and W339 are located close to the sialic acid binding site [24,25], along with residue Y335, which has been shown through mutagenesis (Y335A) to help stabilise sialin in the CO state [25]. The inferred network paths connecting K197 and D350 involve regions close to the sialic acid binding site suggesting a possible functional coupling between the cytosolic ionic lock and the substrate-binding region. In contrast, the R39C mutation reorganises this network in such a way that key hub residues in the wild type lose their central role in the mutant. In the CO conformation, TM1 and residue V211 are no longer important for allosteric communication. In the LO conformation, this reorganisation leads to a loss of the central relay function of both V211 and W339, leading to shorter inferred network paths between K197 and D350 that no longer preferentially involve the region surrounding the sialic acid binding site. These findings suggest that the R39C mutation induces broader dynamical perturbations that may also contribute to the decreased transport activity of sialin in Salla disease.

To assess the computational findings, we mutated a set of 14 residues in the “R39 sphere” to alanine and investigated the effects of these mutations on the lysosomal localisation and transport activity of sialin. Some residues, in particular R39, were also mutated to other amino acids than alanine. These experiments identified four residues, R39 (TM1), E194 (TM5) and E262 (ICH1) of the triplet along with I266 (ICH1), which are critical to preserve a full transport activity and a full delivery of sialin to lysosomes. The essential role of I266 may arise from its role in anchoring ICH1 to the N-terminal bundle to create a hydrophobic cluster that surrounds, and stabilises, the electrostatic interactions of the triplet. Other residues like E264, L267, and S268 among others, for which the models showed little to no crucial interaction, were found to be more dispensable in mutagenesis studies.

A remarkable feature of the mutagenesis experiments is the highly similar outcome of the tested mutations on sialic acid transport and lysosomal localisation (Figure 3A-D), suggesting a common mechanism behind these effects.

In earlier studies, we hypothesised that a global misfolding of R39C sialin could be detected by the quality-control system of the cell, leading defective egress from the reticulum [16,17]. However, this explanation seems unlikely as the sorting mutant of R39C sialin, lacking the dileucine-based (L22/L23) lysosomal sorting motif, as well as other pathogenic missense mutants, are fully delivered to the plasma membrane [1,40]. Based on the similar profiles of the transport data and the lysosomal localisation data in this study, an attractive alternate possibility could be that the recognition of the sorting motif of sialin by adaptor proteins responsible for its intracellular trafficking could be modulated by the conformational state of the protein. As the MD simulations suggest that the R39C and R39K mutations may favor the LO state, we hypothesise that the L22/L23 dileucine motif present in the N-terminus (and unresolved in our models) is less exposed to cytosolic adaptor proteins in the LO state than in the CO state. The rescue of the R39C-induced trafficking defect by an orthosteric ligand which binds sialin in the CO state [16,25] is consistent with this proposal. If this hypothesis holds true, allosteric ligands that counteract the stabilisation of the ionic lock induced by the R39C mutation might simultaneously rescue the lysosomal delivery defect and the transport defect to help treat Salla disease.

## Material and Methods

### Plasmids

Missense mutations were introduced into EGFP-tagged human sialin, sialin R39C, sialin LLGG (L22G/L23G) and sialin LLGG R39C [1] using Q5 Site-Directed Mutagenesis Kit (New England Biolabs). Details of all primers used are provided in Table S1.

### Cell culture and transfection

HeLa and HEK 293 cells were grown at 37°C under 5% CO2 in glucose-rich, Glutamax-I-containing minimum essential medium (MEM) to which 0.1% non-essential amino-acids were added and in glucose-rich, Glutamax-I-containing Dulbecco’s modified Eagle medium (DMEM), respectively, supplemented with 10% foetal bovine serum, 100 U/mL penicillin, and 100 μg/mL streptomycin. HeLa cells were transfected by electroporation using a GHT 1287 electropulsator (Jouan). Typically, 2×106 HeLa cells in 50 μl of ice-cold PBS, pH 7.4, were mixed with 5 μg of plasmid, and immediately subjected to ten square pulses (200 V, 3 ms) delivered at 1 Hz by 4-mm-spaced electrodes. The cells were then diluted with 7 ml of culture medium and distributed in 12 wells (15-mm diameter) of a 24-well culture plate containing glass coverslips. HEK 293 cells were plated (200000 cells/well) into poly-D-lysine-coated 24-well plates and transfected the following day with Lipofectamine 2000 (Invitrogen), according to the manufacturer’s protocol. HeLa and HEK 293 cells were analysed 2 days after transfection.

### Immunofluorescence and image acquisition

HeLa cells were fixed with 4% paraformaldehyde. After quenching with 50 mM NH4Cl and several washes, cells were permeabilised and blocked with 0.05% saponin and 0.2% BSA in PBS buffer containing Ca^2+^ and Mg^2+^. Coverslips were then incubated for at least 1 h with mouse anti-LAMP1 antibodies (H4A3; Developmental Studies Hybridoma Bank) at 0.75 μg/mL in blocking buffer, washed, and incubated with Cy3-conjugated donkey antimouse antibodies (Jackson ImmunoResearch) at 1.4 μg/mL or Alexa Fluor 555 goat antimouse antibodies (Invitrogen) at 2µg/mL in the same buffer. Coverslips were then washed and mounted on glass slides with Fluoromount-G (Electron Microscopy Sciences). Epifluorescence pictures were acquired under a 100× objective lens with a Nikon Eclipse TE-2000 microscope equipped with a CCD camera (Coolsnap). Sialin/LAMP1 colocalisation was quantitated by assessing the spatial correlation between pixel intensities of the green and red channels. Dual-color images were imported into the Fiji version (http://fiji.sc) of ImageJ, and the Substract Background and Coloc 2 plugins were used to calculate Pearson’s correlation coefficients across 10 cells/condition. Statistical analysis was performed using a paired t-test between the means of Pearson’s coefficient on 3 independent experiments.

For HEK cells used in transport assay, localisation of EGFP-tagged mutant sialin was first imaged under the 10x objective with the same microscope. The total fluorescence reflecting the overall expression level of each plasmid was quantitated by assessing the sum of the values of the pixels in at least 3 images of different experiments.

### Transport assay

HEK 293 cells were washed twice with 500 µl of wash buffer (5mM D-glucose, 140mM NaCl, 1mM MgSO_4_, 20mM MOPS, pH 7.0) and incubated for 15 min at room temperature in 200 µl of uptake buffer (5mM D-glucose, 140mM NaCl, 1mM MgSO_4_, 20mM MES, pH 5.0) containing 100µM Neu5Ac (Activate Scientific) and N-acetyl-[6-3H]neuraminic acid (12.5 nM; 0.05 μCi/well) (American Radiolabelled Chemicals). After two brief ice-cold washes and lysis in 200µL of 0.1N NaOH, the cellular radioactivity in the cells was counted by liquid scintillation with a Tri-Carb 4910TR counter (PerkinElmer). Statistical analysis was performed using a paired t-test between the means of the cellular radioactivity on at least 3 independent experiments.

### Conservation analysis

SLC17 family sequences were obtained via BLAST [41]. The human protein sequences of SLC17A1/NPT1 (UniProt ID: Q14916), SLC17A2/NPT3 (UniProt ID: O00624), SLC17A3/NPT4 (UniProt ID: O00476), SLC17A4/NPT5 (UniProt ID: Q9Y2C5), SLC17A5/sialin (UniProt ID: Q9NRA2), SLC17A6/VGLUT2 (UniProt ID: Q9P2U8), SLC17A7/VGLUT1 (UniProt ID: Q9P2U7), SLC17A8/VGLUT3 (UniProt ID: Q8NDX2), and SLC17A9/VNUT (UniProt ID: Q9BYT1) were obtained from UniProt [42] and were used as queries for BLAST searches. For each transporter sequence a total of 5000 results were generated and the sequences were used for further filtering. Sequences containing the descriptors “like”, “low quality protein”, “putative”, “hypothetical”, “partial”, “predicted”, “unnamed”, or “uncharacterized” were removed. The different BLAST results were pooled together and, in the case, where multiple sequences contained the same BLAST accession number, only one was retained to remove duplicates. Only sequences containing the descriptors “Sodium-dependent phosphate transport protein 1”, “Sodium-dependent phosphate transport protein 3”, “Sodium-dependent phosphate transport protein 4”, “small intestine urate exporter”, “sialin”, “Vesicular glutamate transporter 2”, “Vesicular glutamate transporter 1”, “Vesicular glutamate transporter 3”, “Voltage-gated purine nucleotide uniporter”, “SLC17A1”, “SLC17A2”, “SLC17A3”, “SLC17A4”, “SLC17A5”, “SLC17A6”, “SLC17A7”, “SLC17A8”, “SLC17A9”, “Solute carrier family 17”, “NPT1”, “NPT3”, “NPT4”, “VGLUT2”, “VGLUT1”, and “VGLUT3” were kept. The filtered sequences were aligned using FAMSA2 [43]. The aligned sequences were retained only if they shared ≥30% pairwise sequence identity with all nine representative human SLC17A1-9 reference sequences to ensure the sequences belong to the same family. The final alignment contained a total of 7455 sequences. The sequence logos showing the conservation of residues were generated using WebLogo [44].

To investigate the evolutionary conservation of the R39/E194/E262 triplet and the K197/D350 ionic lock, we examined the co-occurrence of these two motifs in the alignment using an in-house Python script. The triplet was deemed conserved if the position equivalent to R39 in sialin contained arginine and the positions equivalent to E194 and E262 contained acidic residues (aspartate or glutamate). The cytosolic ionic lock was deemed conserved if the position equivalent to K197 contained a basic residue (arginine or lysine) and the position equivalent to D350 an acidic residue. Sequences were categorised into four mutually exclusive groups based on the presence or absence of these features: both ionic lock and triplet, ionic lock only, triplet only, or R39 with neither motif. The fraction of sequences in each category was calculated relative to the total alignment size.

### Model construction

Models were generated for the LO and CO conformations using the available cryo-EM structures [25] (PDB ID 8U3F and 8U3D respectively). The experimental structures were used as input templates for the online server CollabFold [45] employing AlphaFold2 [26] along with the sequence of sialin extracted from UniProt (UniProt ID Q9NRA2) to generate five models of sialin based on each template. The generation of these models was done using default parameters. The top model from each run corresponded in state to the input cryo-EM structure and was used as a template to model the missing loops in the experimental structures, while keeping the existing coordinates unchanged. Using MODELLER [46] the missing segments for the LO (26-34, 69-80, 87-95, and 487-495) and CO (26-36, 70-97, and 488-495) states were modelled. Residue R168 in the LO experimental structure (PDB ID 8U3F) was turned into a lysine to be in agreement with the wild-type sequence. A total of 1000 models were generated for each state and the best one was selected based on the DOPE score [27]. The best model for each state was used as the starting structure for MD simulations.

### Selection of protonation state

The residue protonation states were selected in a two-step process involving the wild-type models of the LO and CO conformations. Initially, PROPKA3 [47] was run on the models of the LO and CO conformation of sialin. Residues that are located on the luminal side of the transporter and whose p*K*_a_ was predicted to be >4.5 were protonated. The residues E75, E85, H86, H93, H94, E106, E326, E455 were initially protonated in both conformations. MD simulations were run for 450 ns in triplicate for each state and TrIPP [48] was run on the trajectories to determine the p*K*_a_ values of titratable residues during the trajectory. Residues that showed a p*K*_a_ value >4.5 for >50% of the trajectory in all three replicas and were in the luminal side of the protein were selected for protonation. The initial models of the transporter were reprotonated at residues E75, H86, H93, H94, E106, E326, D388, and E455 in both states. Residue E85 was additionally protonated in the LO but not in the CO state. MD simulations were then run for 450 ns in triplicate. The protonation state of the mutant (R39C and R39K) was the same as in the wild type depending on the state.

### System Setup and Simulation Protocol

The generated protein structures were embedded in a model membrane of 1-palmitoyl-2-oleoyl-sn-glycero-3-phosphocholine (POPC) lipids using the CHARMM-GUI Membrane builder [49,50]. The PPM 2.0 server [51] was used to orient the protein in the membrane. A rectangular box was used, with water thickness (minimum height of water above and below the system) of 22.5 Å and an initial XY length of 90 Å, resulting in a number of POPC lipids for the lower and upper leaflets of ∼ 94-96. The system was solvated with TIP3P molecules. Protons were added to the residues that were selected to be protonated (see Selection of protonation states). The coordinates of the protons were extracted from the pqr file generated by running the online PROPKA3 [47] server. The mutants R39C and R39K were generated by using the Build Mutant tool in Discovery Studio 2025. All coordinates of the system, except the ones of the mutated residue were kept the same. The GROMACS command *gmx genion* was used to add counterions to neutralise the system.

MD simulations were performed with GROMACS 2023.1. Periodic boundary conditions were applied, and a rectangular box was used. The protein, lipids and ions were described using CHARMM36m [52]. The Particle Mesh Ewald (PME) method [53] was used for electrostatic interactions with a cutoff of 12 Å for direct-space sum. A 12-Å cutoff was also used for van-der-Waals interactions. The equations of motion were integrated using the leap-frog method with a 1-fs time step for most of the equilibration steps and 2-fs for the last equilibration step and the production run. LINCS [54] was used to constrain bonds involving hydrogen atoms.

Energy minimisation, equilibration and production were run as described in Matsingos et al. 2024 [55]. Briefly, the system was energy-minimised using 50000-steps of steepest descent. Equilibration was carried out in different steps as described in Table S2. Positional restraints were first applied on both the protein and lipids heavy atoms and gradually decreased while equilibrating the temperature to 300 K and the pressure to 1 bar. The Berendsen thermostat [56] was used for the initial stage and subsequently replaced with Nosé-Hoover [57]. For the pressure coupling, the semi-isotropic Berendsen barostat [56] was initially used and replaced by the Parrinello-Rahman [58] for the last NPT stages. Production NPT runs were performed for 450 ns for all simulations. Three equilibration + production replicas were run for all mutants (wild type, R39C, and R39K,) and both the LO and CO conformations.

### Trajectory analyses

Analyses were carried out on the last 300 ns of the production run of the trajectories sampled every 100 ps.

Distances, hydrogen bonds, and bundle angle were measured using MDAnalysis 2.10.0 [59,60]. Hydrogen bonds were determined to be present if the distance between the donor (D) and acceptor (A) was <3.5 Å and the angle A-D-H was <30°. The RMSF was calculated using the *gmx rmsf* command of GROMACS.

To analyse the allosteric communication in the trajectories, dynamic cross-correlation (DCC) matrices were calculated using the dccm function of the bio3d [61] R library on the last 300 ns of the production run of each replica for each mutant and state. As in previous work [55,62,63], A consensus DCC matrix was constructed by averaging correlation values across the three replicas and only keeping the correlations |r|>0.3 in all three replicas. This was done to focus on correlations consistent across replicas, minimising noise from inter-replica variability, and false positive couplings, and providing correlations that may be more related to the native protein behaviour rather than the particular conformational space sampled by each replica. The cutoff of 0.3 was chosen since it allowed for >99% network connectivity across all conditions, preserving the integrity of the allosteric path analysis. A correlation network was generated for each mutant and state where the Cα atoms of the protein represent nodes in the network connected by edges representing correlations |r|>0. For each network, the betweenness centrality was calculated and normalised to a range between 0 and 1. Optimal and suboptimal path analysis was also performed between the residues of the cytosolic ionic lock K197 and D350 using the *cnapath* function to identify the top 500 shortest paths.

Non-bonded interactions and d_1_ to d_4_ distances were analysed using the *Analyze Trajectory* protocol implemented in Discovery Studio Simulation 2025. GROMACS trajectory files were converted into DCD format prior to analysis. Hydrogen bonds, hydrophobic contacts, and electrostatic interactions were considered, as defined by the default distance- and angle-based geometric criteria implemented in Discovery Studio. The resulting interaction data were processed and analysed using custom Python scripts. For each mutant and conformational state, only the final 300 ns of the production phase of each replica were included in the analysis.

## Credit authorship contribution statement

C.M and I.L. contributed equally as first authors. C.M. performed the modelling experiments under the supervision of F.A. and A.G.L. while I.L. conducted the cellular assays under the supervision of C.A.. M.B. and A.G.L. contributed to the analysis of the modelling data. M.V., J.M., C.D. participated in the cellular assays. C.A., B.G. and F.A. initiated the project. C. M. drafted the manuscript with contributions from F.A., C.A. B.G. and the other authors. All authors approved the final version of the manuscript.

## Data availabilty

Model coordinates and plasmids are available upon request.

## Acknowledgements

CM has been funded by the Fondation Dassault Systèmes [Grant number: 287103, 05/05/24-12/20/26]. The work was supported by grants from the Salla Treatment and Research (STAR) Foundation [07/01/23-12/31/24], including a postdoctoral fellowship to I.L., and from the non-profit organisation “Vaincre les Maladies Lysosomales (VML)” [12/01/23-12/01/25]. The Science Ambassador Program from Dassault Systèmes BIOVIA is acknowledged for the support of F.C.A. This project was provided with computing HPC and storage resources by GENCI at CINES thanks to the grant 2024-AD010715831 on the supercomputer Adastra’s MI250 partition. Some image acquisition were performed at the SCM facility (BioMedTech Facilities, INSERM US36| CNRS UAR2009 | Université Paris Cité, https://biomedicale.u-paris.fr/biomedtech-facilities/). DNA dosage and quantification of total fluorescence of HEK cells on plates were performed at the Cyto2BM facility (BioMedTech Facilities, INSERM US36| CNRS UAR2009 | Université Paris Cité, https://biomedicale.u-paris.fr/biomedtech-facilities/).

