## supplement for "Molecular basis of Salla Disease: R39C Mutation Effects on the Lysosomal Transporter Sialin"

**[a]. Université Paris Cité, CNRS, Saints-Pères Paris Institute for the Neurosciences, F-75006 Paris, France**
**[b]. Biovia, Dassault Systèmes, F-78140 Vélizy-Villacoublay, France**
**[c]. Biovia Science Council, Dassault Systèmes, F-78140 Vélizy-Villacoublay, France**

**[#] these authors contributed equally**

**Correspondence to Christine Anne, Francine C. Acher, and Christos Matsingos :** (*C. Anne*), (*F. Acher*), (*C. Matsingos*)

### Table of Contents

**Figure S1:** . Interactions of R39 in the Lumen- and Cytosol-open states of Sialin ..........................................**S3**
**Figure S2:** Solvent accessible surface around R39 ..........................................**S5**
**Figure S3:** Effect of mutations on the lysosomal localisation of sialin ..........................................**S7**
**Figure S4:** The R39C, R39K, E194A, E262A and I266A mutations do not impair surface expression of GFP-sialin L22G L23G in HEK cells ..........................................**S8**
**Figure S5:** Mutations around the Sialin R39 site do not impair surface expression of GFP-Sialin L22G L23G in HEK cells..........................................**S9**
**Figure S6:** Effect of mutations on the total cellular expression of human L22G L23G Sialin..........................**S10**
**Figure S7:** Violin plots showing the distances between the heavy atoms of R39 and its surrounding residues ..........................................**S11**
**Figure S8:** Wild-type interactions in MD simulations in the LO and CO conformations at R39 site..S12........**S12**
**Figure S9:** Non-bonded interaction monitoring of the wild-type and R39C CO trajectories..........................**S13**
**Figure S10:** Hydrophobic interaction monitoring of the wild-type and R39C LO trajectories........................**S14**
**Figure S11:** Hydrophobic interaction monitoring of the wild-type and R39C CO trajectories........................**S15**

**Figure S12:** Distances $d_1$ to $d_4$ of triplet, relay and ionic lock interactions in WT sialin across three MD simulation replicas……**S16**
**Figure S13:** Distances $d_1$ to $d_4$ of triplet, relay and ionic lock interactions in WT sialin across the three MD simulation replicas in the CO conformation……**S17**
**Figure S14:** Differences in R39 site interactions between R39C mutant and wild-type sialin in the CO conformation……**S18**
**Figure S15:** Hydrogen bonding interactions of E194 and E262……**S19**
**Figure S16:** Line plots showing the evolution of the R39-E262 Cα distance……**S20**
**Figure S17:** Line plots showing the RMSF values of the Cα atoms of the wild-type and R39C mutant sialin during MD simulations of the LO and CO conformations……**S21**
**Figure S18:** R39K mutant interactions in the LO conformation……**S22**
**Figure S19:** R39K mutant interactions in the CO conformation……**S24**
**Figure S20:** Non-bonded interaction monitoring of the wild-type and R39K LO trajectories……**S26**
**Figure S21:** Non-bonded interaction monitoring of the wild-type and R39K CO trajectories……**S27**
**Figure S22:** Histograms showing the distribution of dynamic cross-correlation (DCC) values between wild-type (WT) and R39C sialin mutant……**S28**
**Figure S23:** Bundle angle in the LO conformation of wild-type and R39C mutant sialin……**S29**
**Figure S24:** Schematic representation of network analysis……**S30**

**Table S1:** Primers used for site-directed mutagenesis……**S31**
**Table S2:** Sequence of equilibration steps for the MD simulations……**S33**

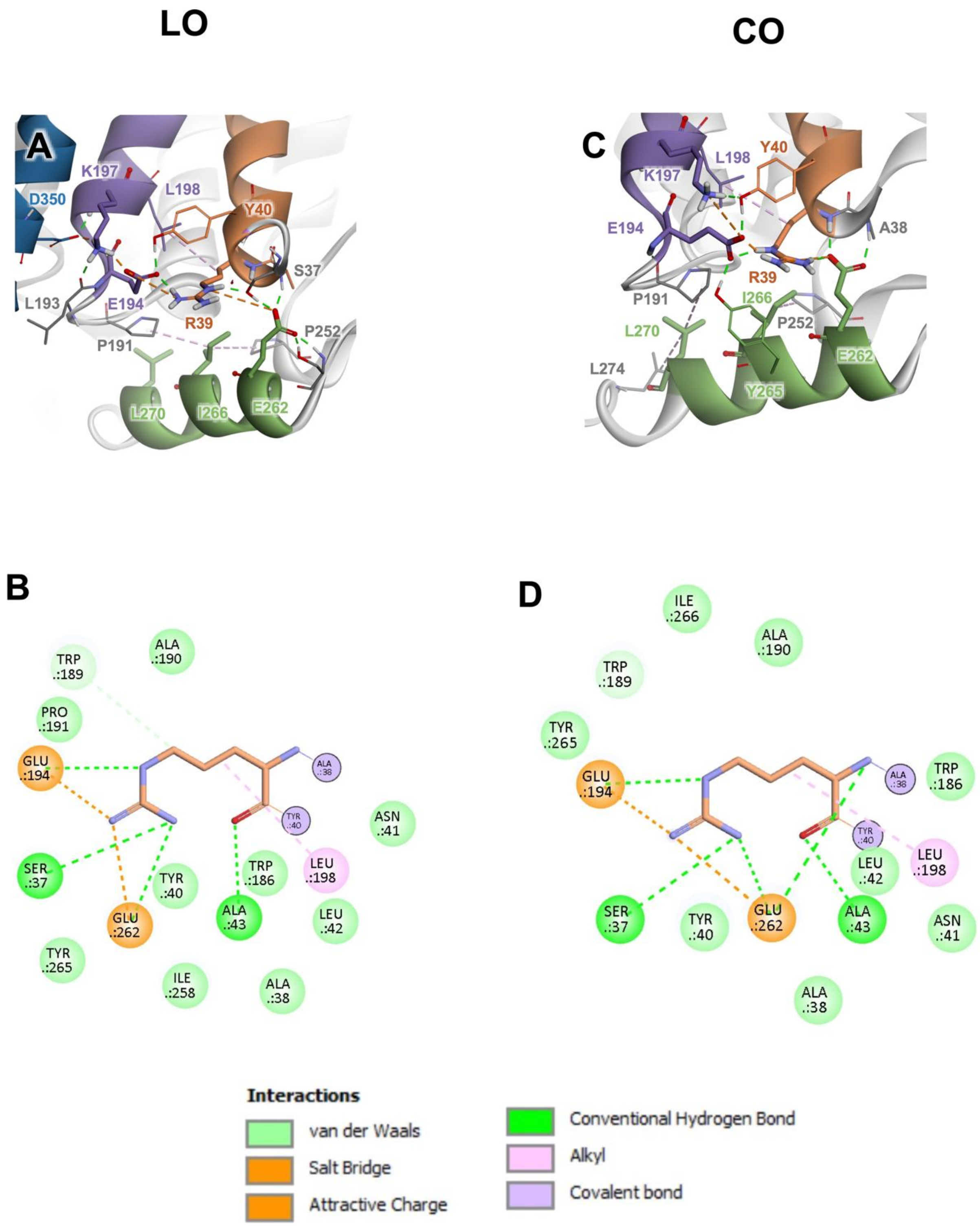


**Figure S1.** Interactions of R39 in the Lumen- and Cytosol-open states of Sialin. The interactions of R39 in the LO (A-B) and the CO (C-D) states are shown. Residue R39 is shown to interact with E194 and E262 over hydrogen bonds (green dashed line) and salt bridges (orange dashed line) in both the LO (A-B) and CO (C-D) conformations. Residue R39 is very tightly packed in both the LO (A-B) and CO (C-D) conformations forming apolar interactions (purple dashed line) with L198 in both states (B and D) and van-der Walls contacts with A31, A32, V34, S37, Y40, N41, L42, A43, W186, A190, P191, W186, E194, L198, I 258, E262, Y265, and I266.

A schematic 2D-diagram of the interactions in the LO (C) and CO (F) states is also shown with salt bridges (orange bashed lines), hydrogen bonds (green dashed lines), apolar interactions (purple dashed lines) and van-der-Waals contacts (light green circles).

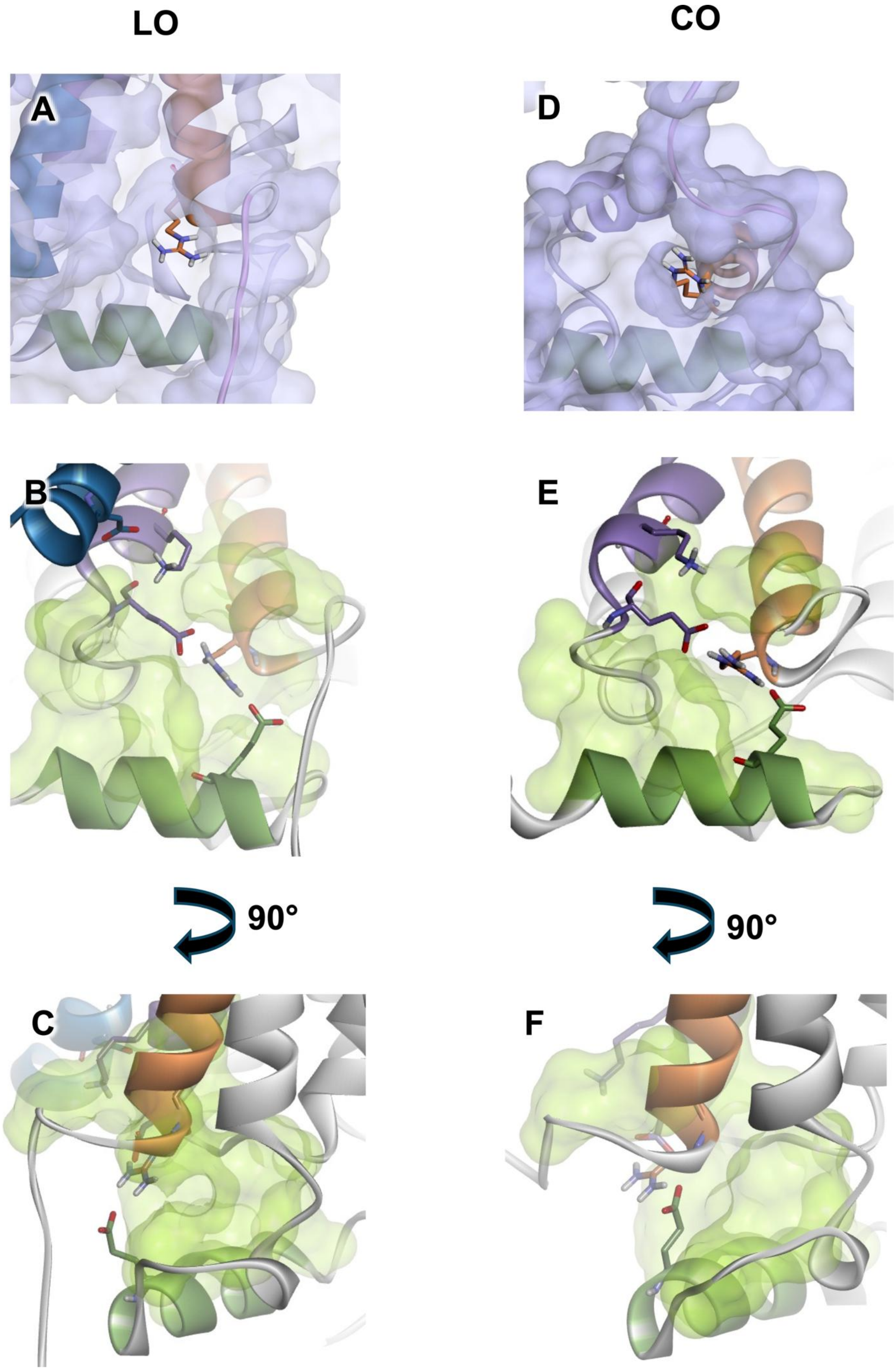
LO
CO
A
D
B
E
90°
90°
C
F

**Figure S2.** A and D: Solvent accessible surface (blue) around R39 shown for the LO (A) and CO (D) conformations. B-C, E-F: Solvent accessible surface (green) of the hydrophobic cluster around R39 in LO and CO conformations (V34, A38, Y40, A43, W189, A190, P191, P192, L193, L198, P252, I258, Y265, I266, L270). B and D Front view, C and F side view.

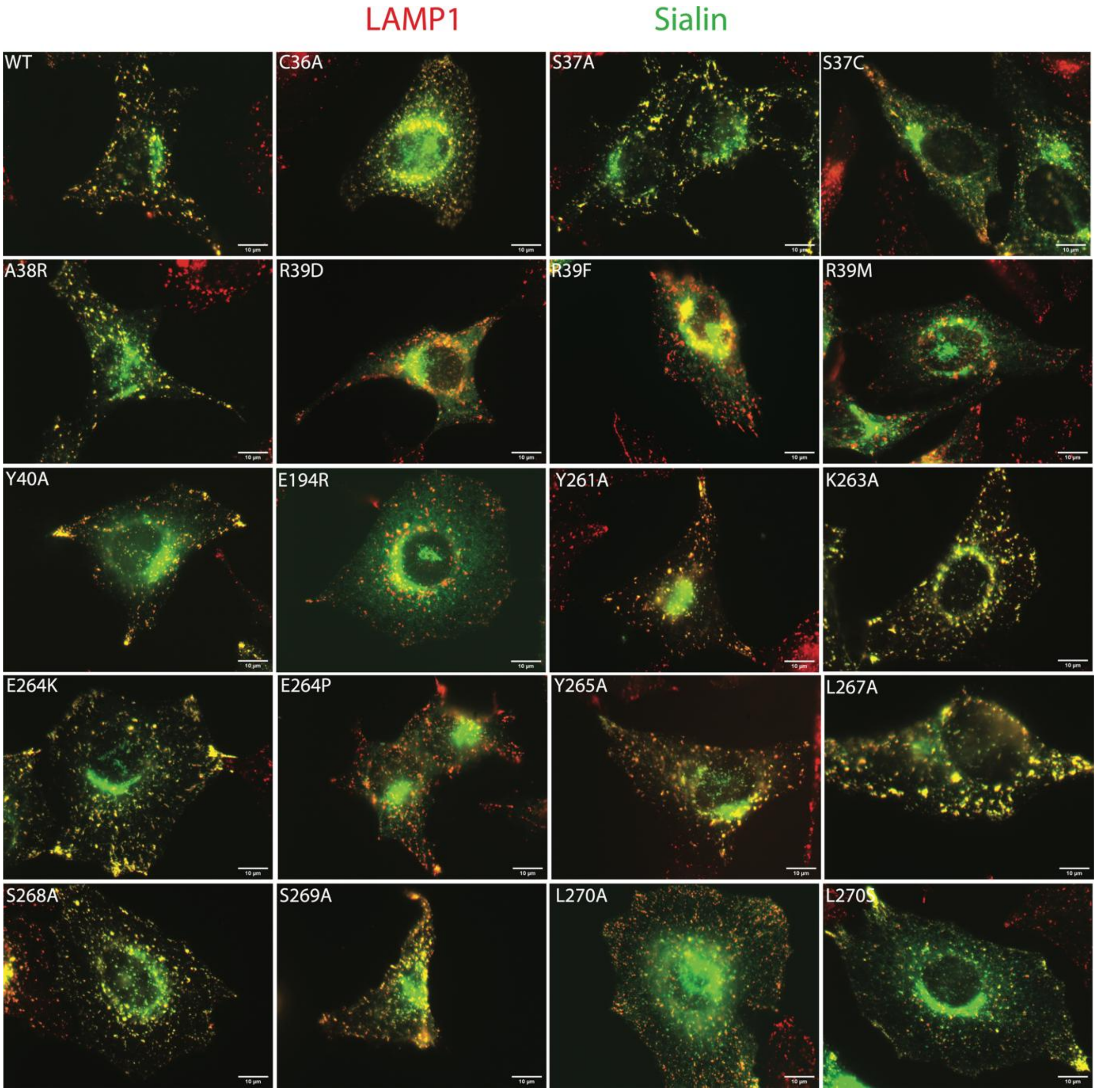


**Figure S3.** Effect of mutations on the lysosomal localisation of sialin. Different mutated human sialin tagged with EGFP (green) constructs were transiently expressed in HeLa cells by electroporation. After two days, cells were fixed and analysed under fluorescence microscopy using LAMP1 immunostaining (red) to detect late endosomes and lysosomes. Only the merges of the photos in red and green are shown here. The scale bar represents 10 µm.

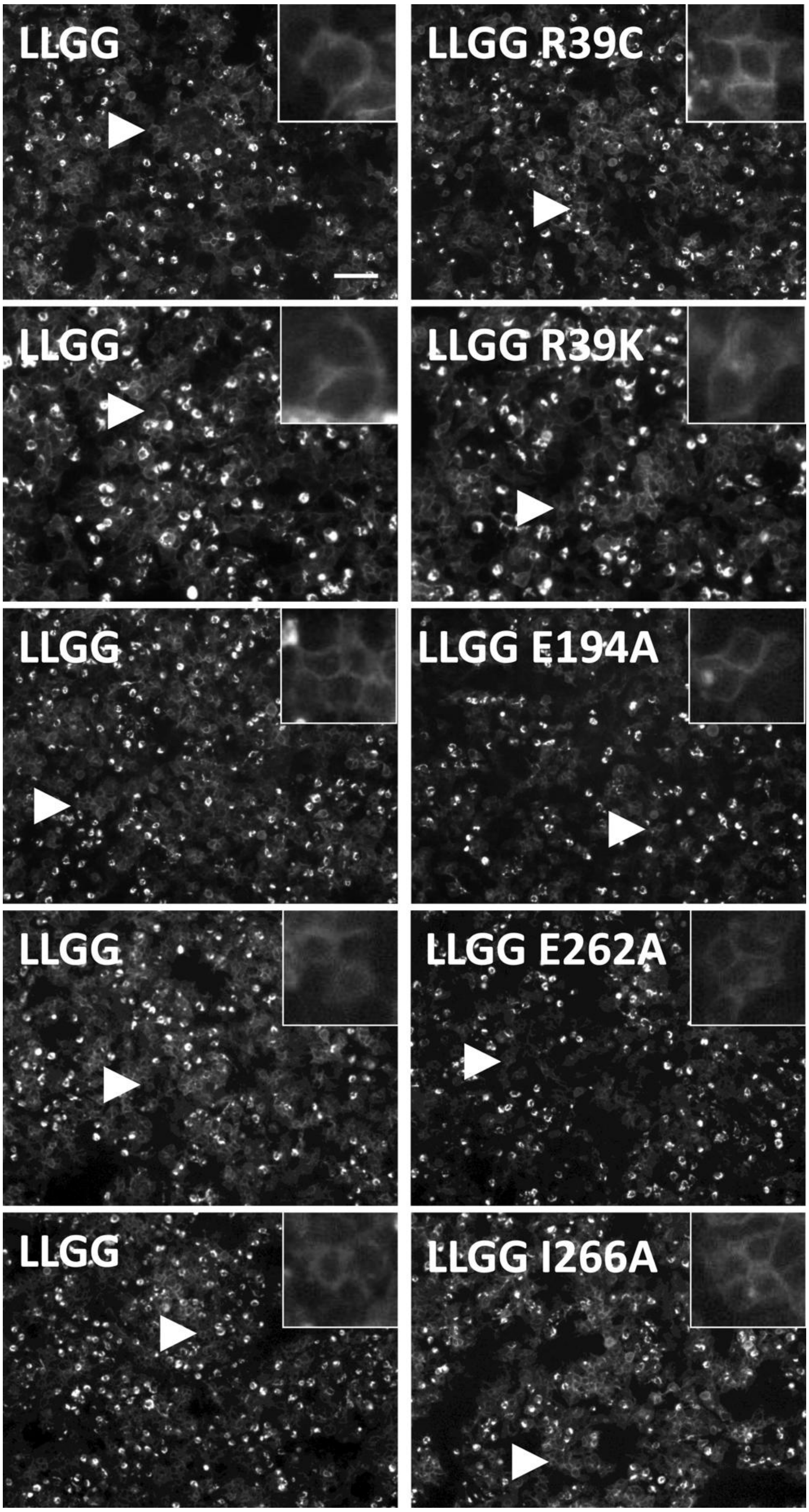


**Figure S4.** The R39C, R39K, E194A, E262A and I266A mutations do not impair surface expression of GFP-sialin L22G L23G in HEK cells. EGFP-tagged human L22G L23G sialin, without (control) and with additional mutations, was transiently transfected by lipofection in HEK cells. After two days, cells were observed under fluorescence microscopy to control constructs' intracellular distribution before the transport assay. Insets are a magnification of the cells indicated by the arrowheads. The scale bar represents 100 µm.

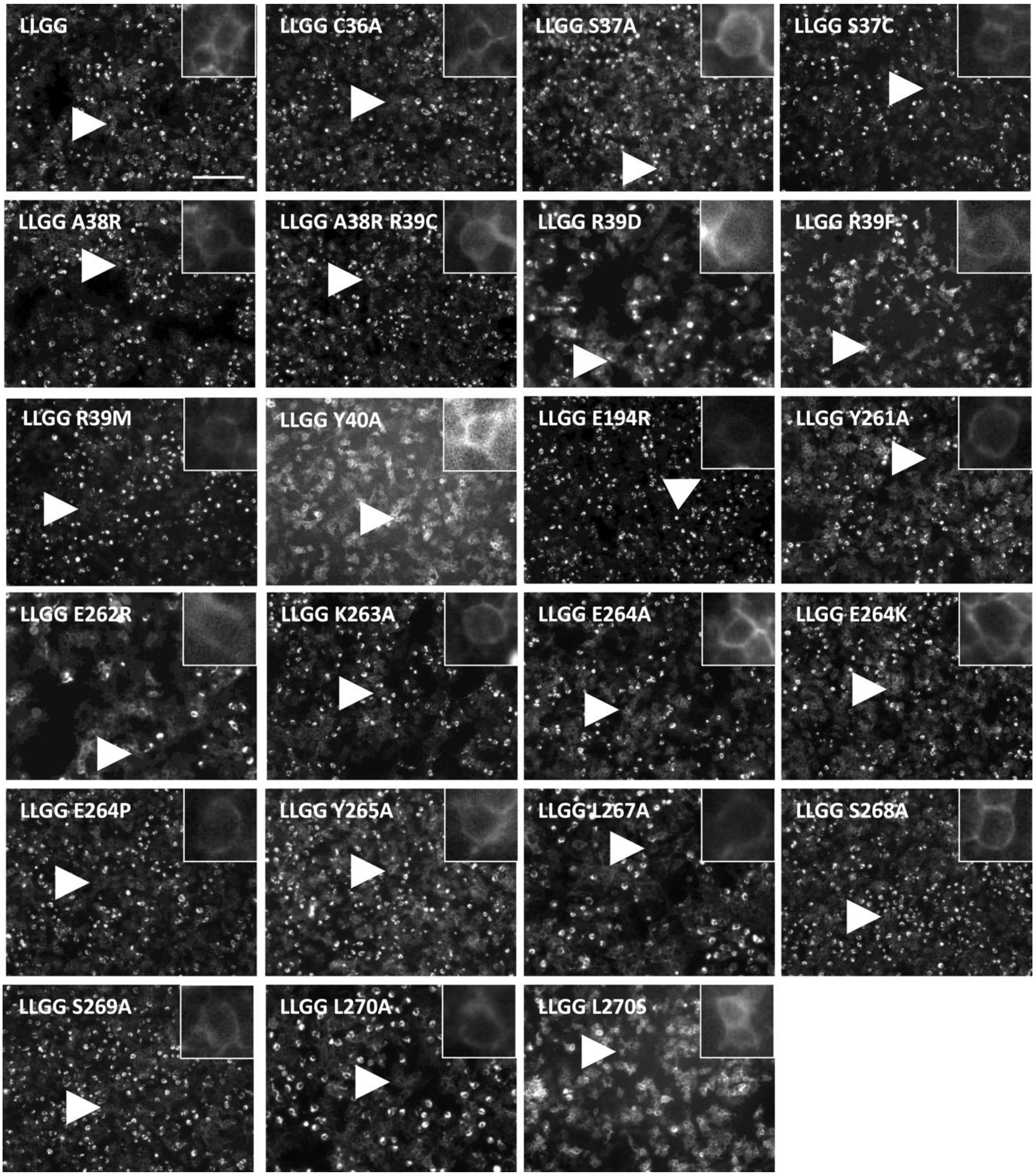


**Figure S5.** Mutations around the Sialin R39 site do not impair surface expression of GFP-Sialin L22G L23G in HEK cells. EGFP-tagged human L22G L23G sialin without (control) and with additional mutations were transiently transfected by lipofection in HEK cells. After two days, cells were observed under fluorescence microscopy to control constructs' intracellular distribution before transport assay. Insets are magnification of the cells indicated by the arrowheads. The scale bar represents 100 µm.

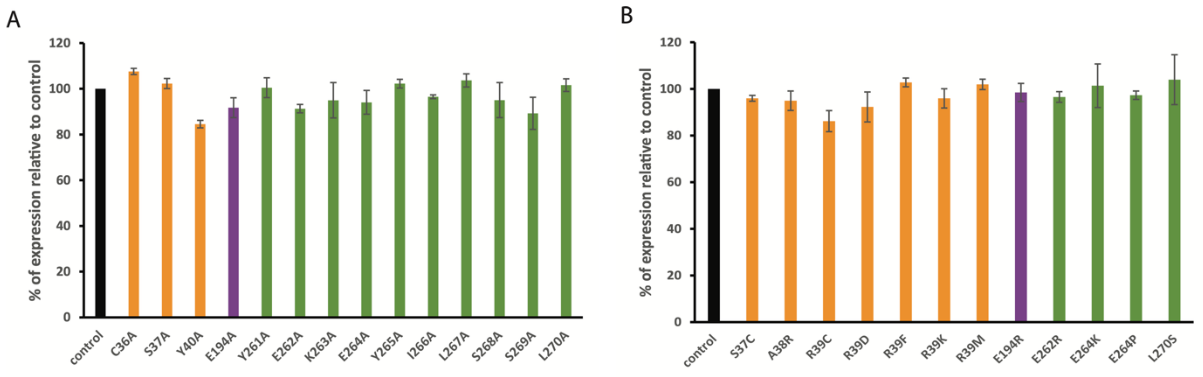


**Figure S6.** Effect of mutations on the total cellular expression of human L22G L23G Sialin. Human L22G L23G Sialin tagged with EGFP with/without mutations were transiently transfected by lipofection in HEK 293T cells. Before the whole-cell transports assays, the levels of sialin expression was analysed under fluorescence microscopy. The sum of the values of the pixels in the large field images was measured and the values for the mutants were normalised to that of control sialin in each experiment.

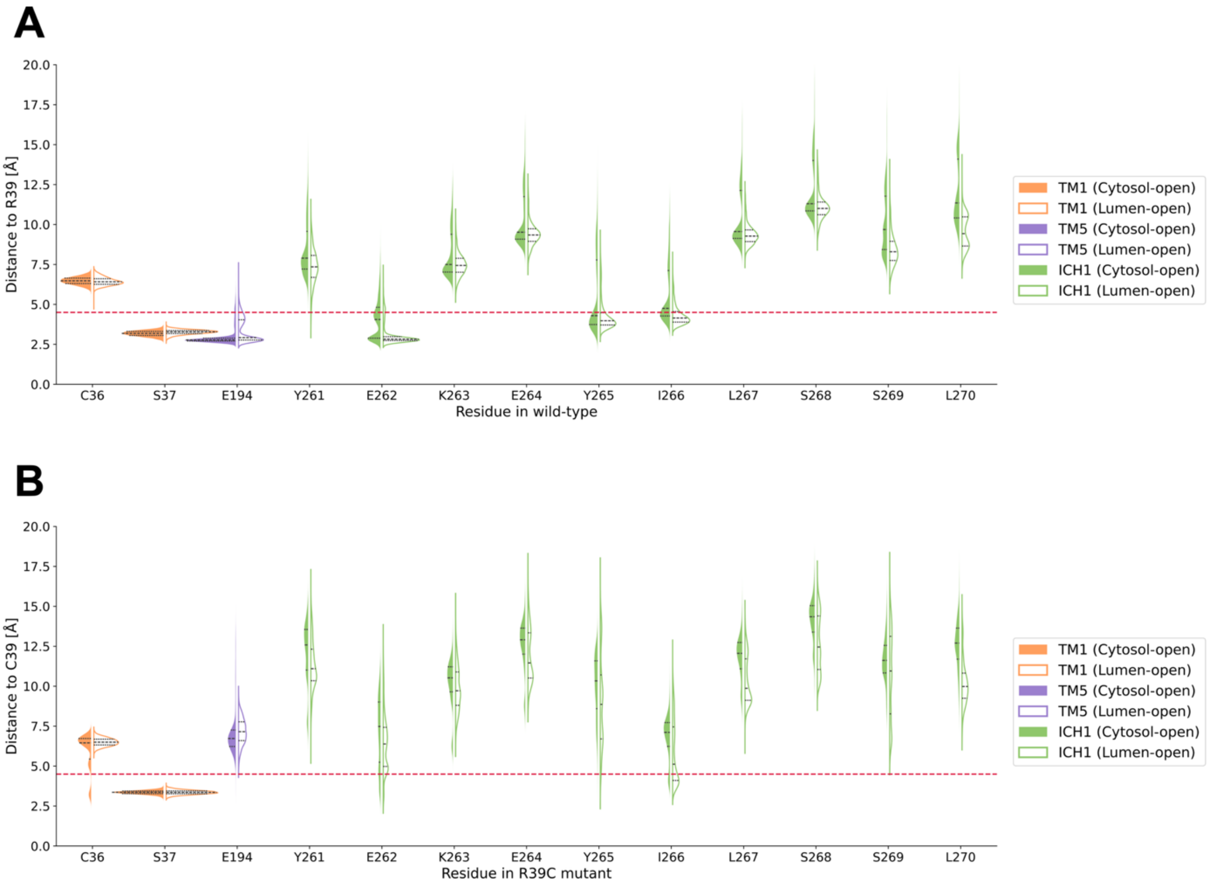


**Figure S7.** Violin plots showing the distances between the heavy atoms of R39 and its surrounding residues. Residues belonging to TM1 (orange), TM5 (purple), and ICH1 (green) are shown and distance calculated in the CO (filled in left violin plots) and LO (not filled in right violin plots) conformations. The distances are shown for wild type (A) and R39C mutant (B) sialin. Residues with a minimum heavy atom distance <4.5 Å (red dashed line) are considered in contact. The data from the last 300 ns of all three replicas (MD1-MD3) are shown.

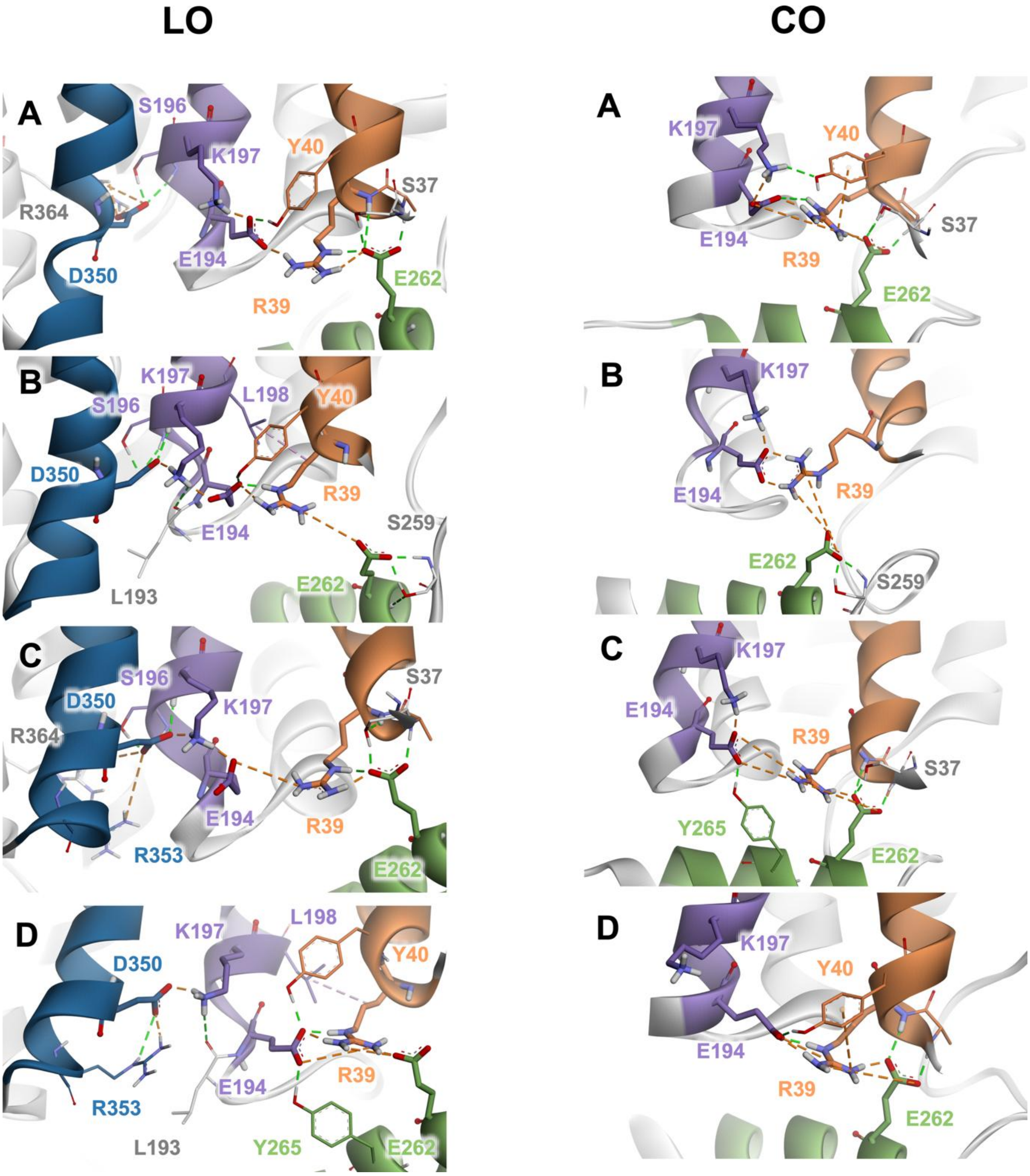


**Figure S8.** Wild-type interactions in MD simulations in the LO and CO conformations at R39 site. In wild-type sialin, the R39 site adopts four distinct configurations during the simulations: A) E194 forms strong interactions with both R39 and K197, while R39 simultaneously interacts strongly with E262; B) E194 maintains strong interactions with R39 and K197, R39 moves away from E262 but remains within interaction range, and K197 interacts with D350 (in the LO conformation); C) E194 interacts strongly with K197 and is distant from R39 but still within interaction range, R39 forms a strong interaction with E262, and K197 interacts with D350 (in the LO conformation); and D) R39, E194, and E262 form a strongly interacting triplet, E194 does not interact with K197, and K197 interacts with D350 (in the LO conformation).

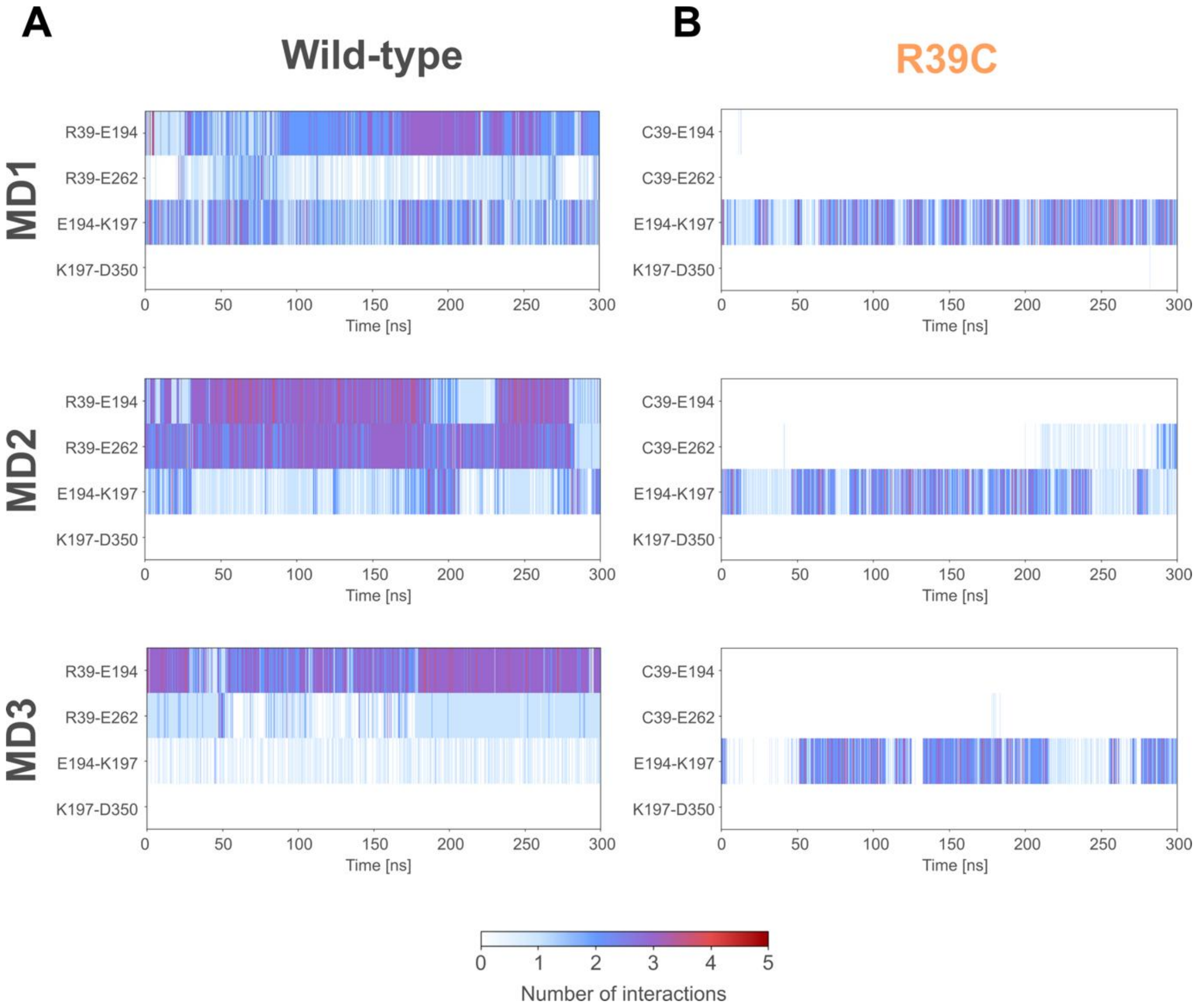


**Figure S9.** Non-bonded interaction monitoring of the wild-type (A) and R39C (B) CO trajectories. The interactions for the C/R39-E194, C/R39-E262, E194-K197, and K197-D350 pairs are shown during MD simulations for wild-type (A) and R39C mutant (B) sialin in the CO conformation. The data of the last 300 ns of the production run of all three replicas, MD1 (top), MD2 (middle), and MD3 (bottom), are shown for each mutant. Interactions are shown as vertical lines, where the colour of the line indicates the number of interactions, ranging from 0 (white) to 5 (red).

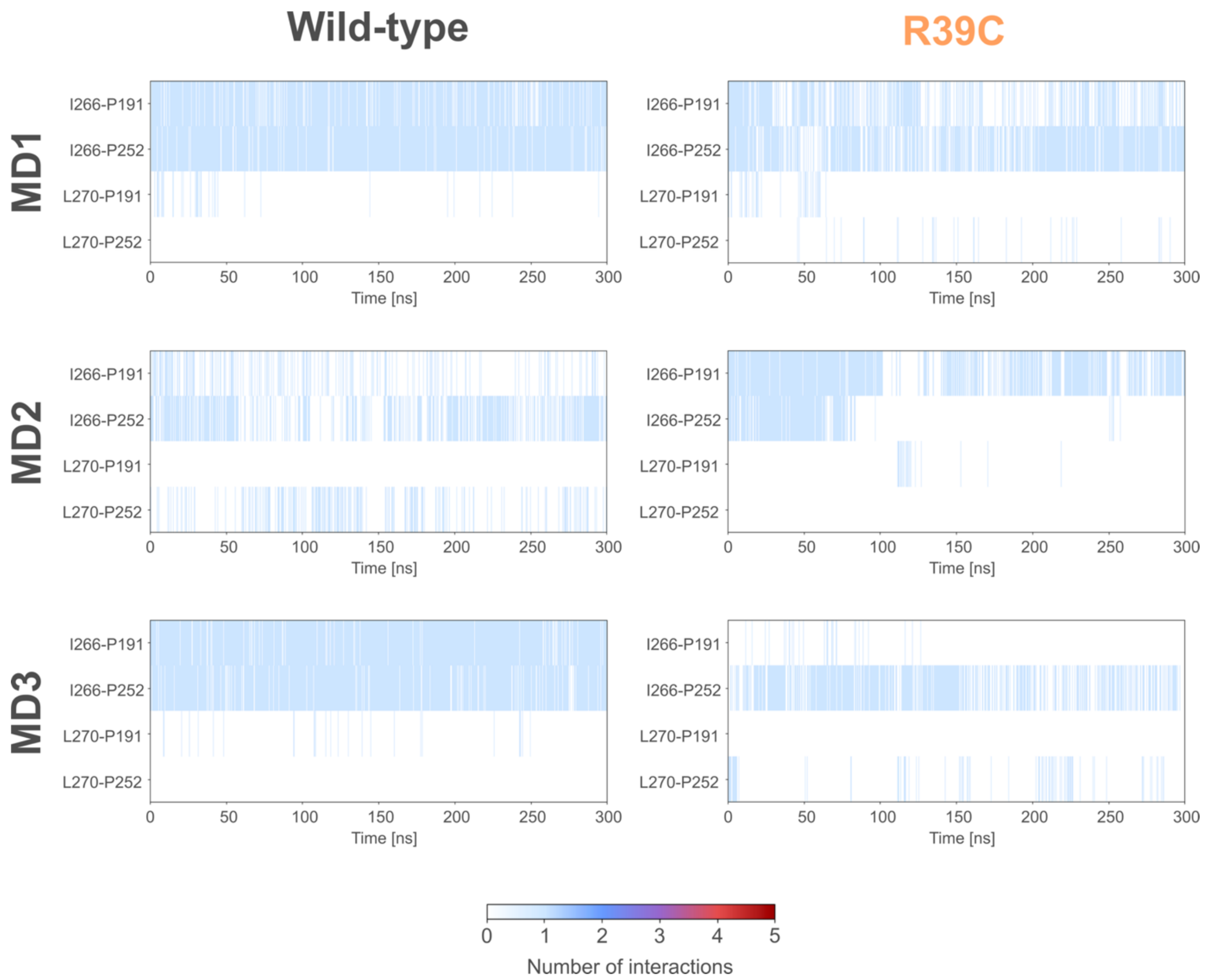


**Figure S10.** Hydrophobic interaction monitoring of the wild-type and R39C LO trajectories. The interactions for the I266-P191, I266-P252, L270-P191, and L270-P252 pairs are shown during MD simulations for wild-type (left) and R39C mutant (right) sialin in the LO conformation. The data of the last 300 ns of the production run of all three replicas, MD1 (top), MD2 (middle), and MD3 (bottom), are shown for each mutant. Interactions are shown as vertical lines, where the colour of the line indicates the number of interactions, ranging from 0 (white) to 5 (red).

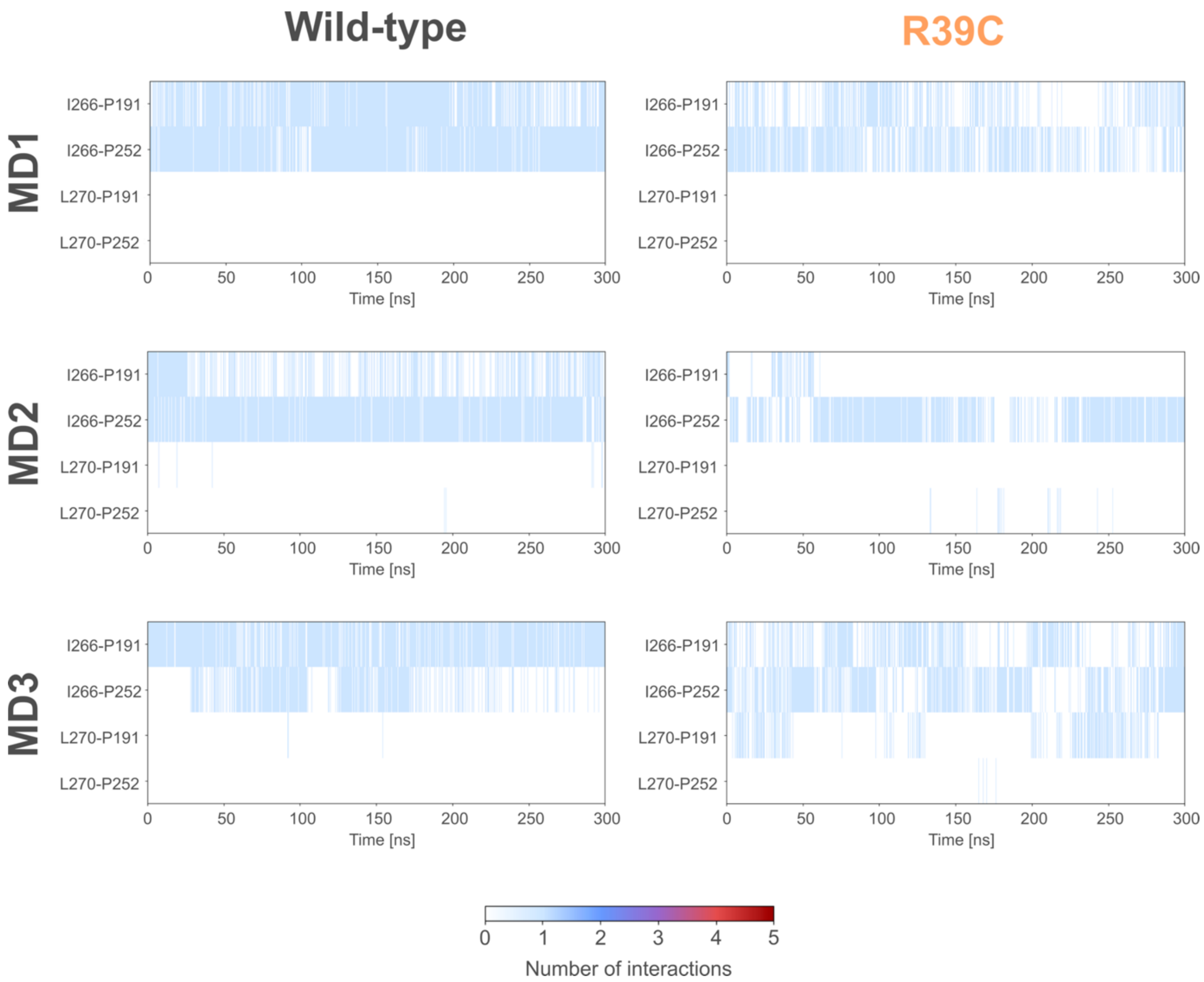


**Figure S11.** Hydrophobic interaction monitoring of the wild-type and R39C CO trajectories. The interactions for the I266-P191, I266-P252, L270-P191, and L270-P252 pairs are shown during MD simulations for wild-type (left) and R39C mutant (right) sialin in the CO conformation. The data of the last 300 ns of the production run of all three replicas, MD1 (top), MD2 (middle), and MD3 (bottom), are shown for each mutant. Interactions are shown as vertical lines, where the colour of the line indicates the number of interactions, ranging from 0 (white) to 5 (red).

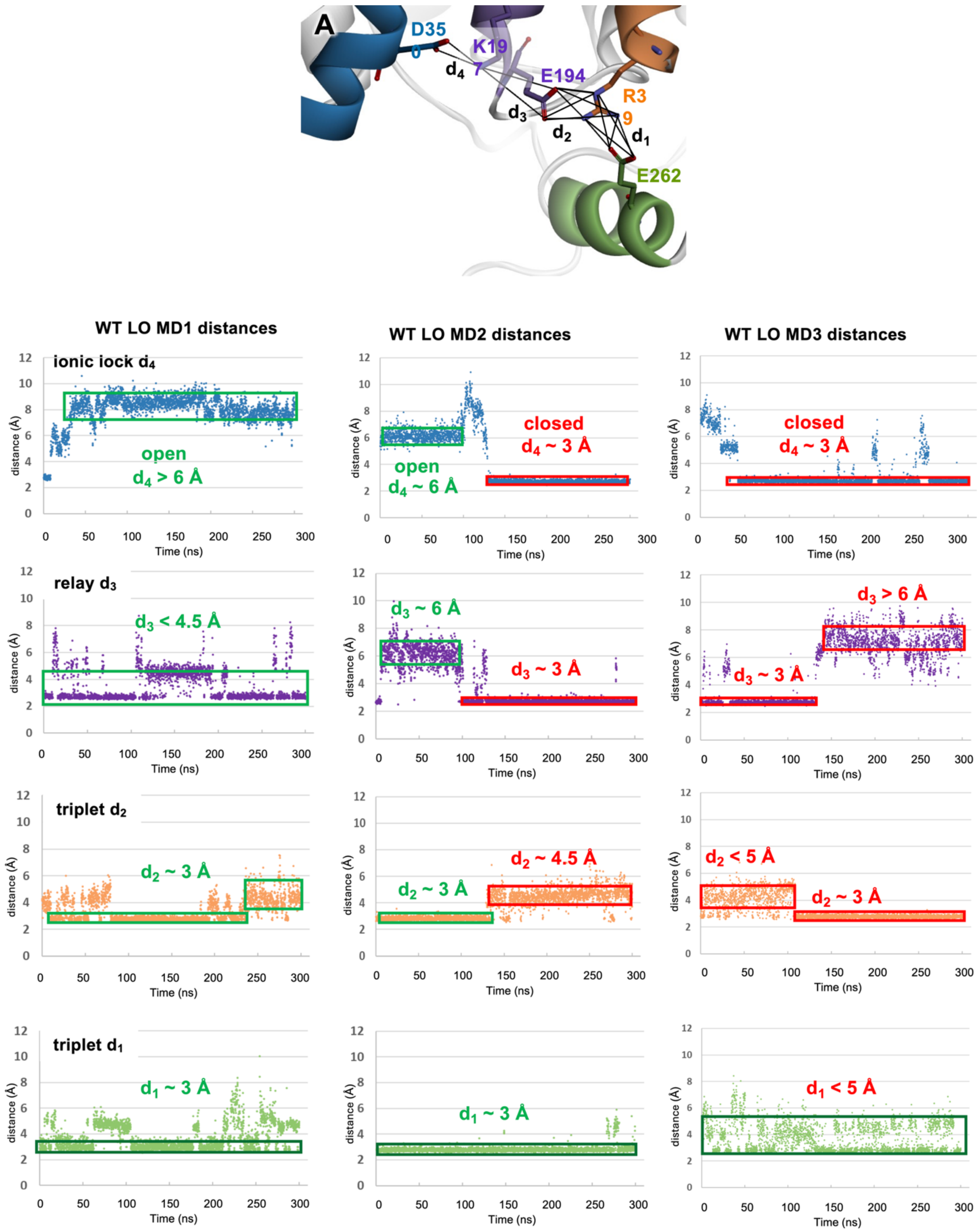


**Figure S12.** Distances $d_1$ to $d_4$ of triplet, relay and ionic lock interactions in WT sialin across three MD simulation replicas (data from the last 300 ns of all three replicas MD1-MD3) in the LO conformation. The ionic lock is open when the relay and triplet make strong interactions (d < 4.5 Å) as in MD1 or when the relay is weak ($d_2$ ~ 6 Å) but still able to transmit the effect of strong triplet interactions ($d_2$ ~ 3 Å) as in MD2. The ionic lock is closed when the relay is strong ($d_3$ ~ 3 Å) but the triplet too weak ($d_2$ > 4.5 Å) as in MD2 or when the relay is impaired ($d_3$ > 6 Å) even if the triplet interaction is strong ($d_2$ ~ 3 Å) as in MD3.

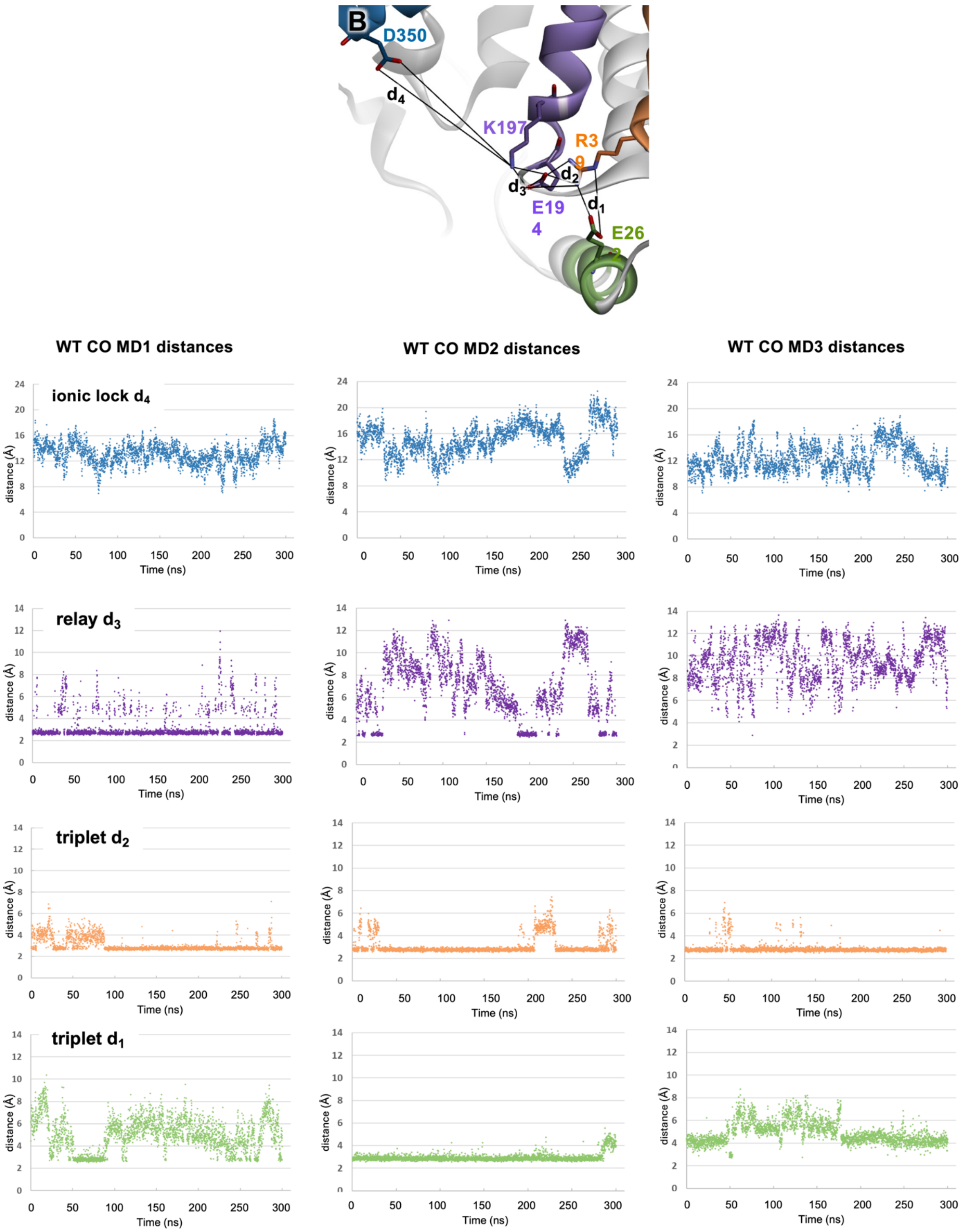


**Figure S13.** Distances $d_1$ to $d_4$ of triplet, relay and ionic lock interactions in WT sialin across the three MD simulation replicas in the CO conformation (data from the last 300 ns of all three replicas MD1-MD3). While the relay interaction ($d_3$ from 3 to 12 Å) is quite variable, the triplet interactions remain strong (d ~ 3 Å).

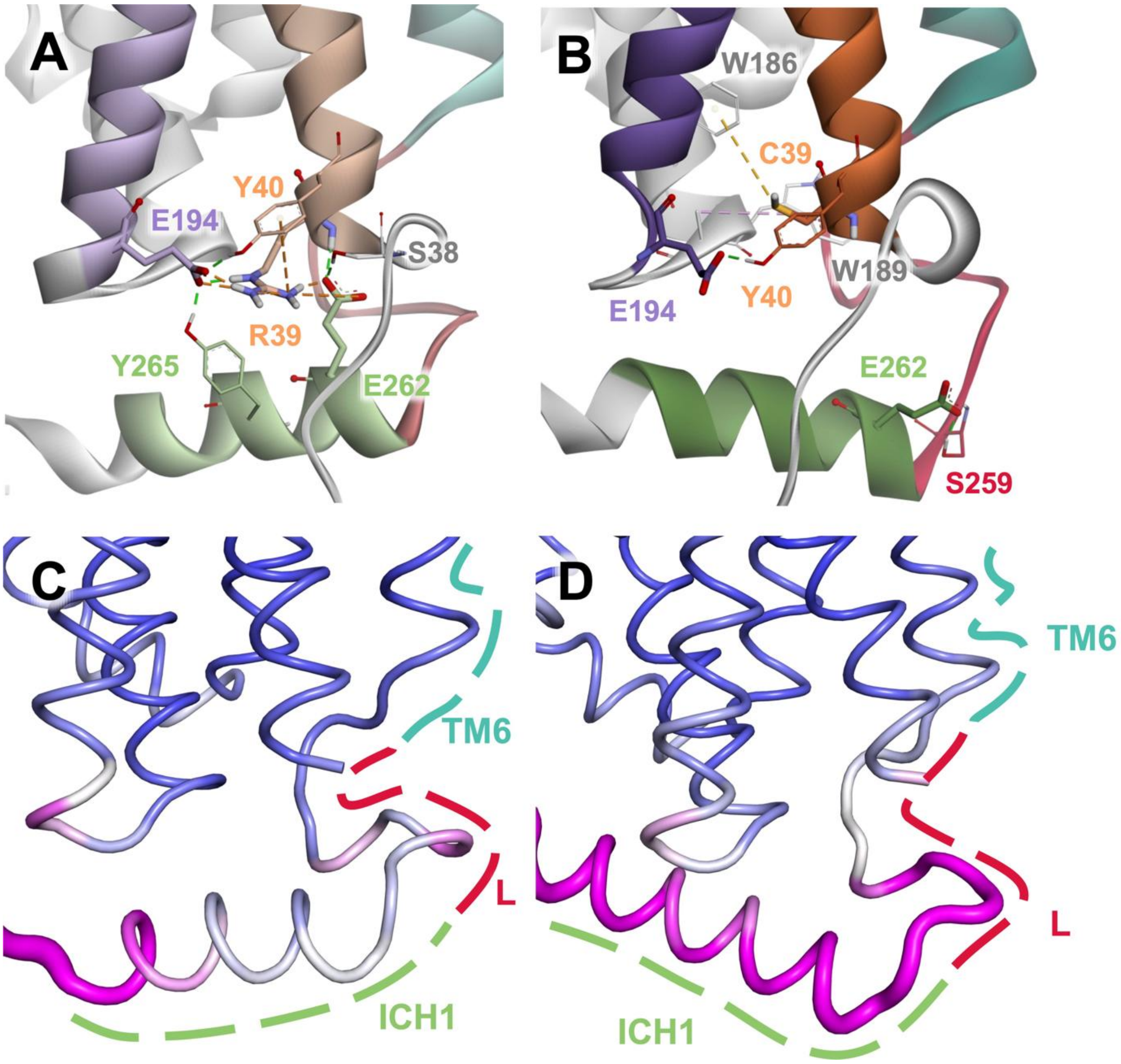


**Figure S14.** Differences in R39 site interactions between R39C mutant and wild-type sialin in the CO conformation. A and B) Interactions of the R39 site in wild-type (A) and R39C mutant (B) shown on representative structures of the CO conformation. In the wild-type, R39 is able to form hydrogen bonds (dashed green lines) and salt bridges (orange dashed lines) with E194 and E262, while in the R39C mutant, these interactions are not possible. C and D) The RMSF of the Cα atoms is shown mapped on the average structures of wild-type (C, MD2) and R39C mutant (D, MD1) sialin LO trajectories. High RMSF values are shown as thick and magenta, and small ones as thin and blue.

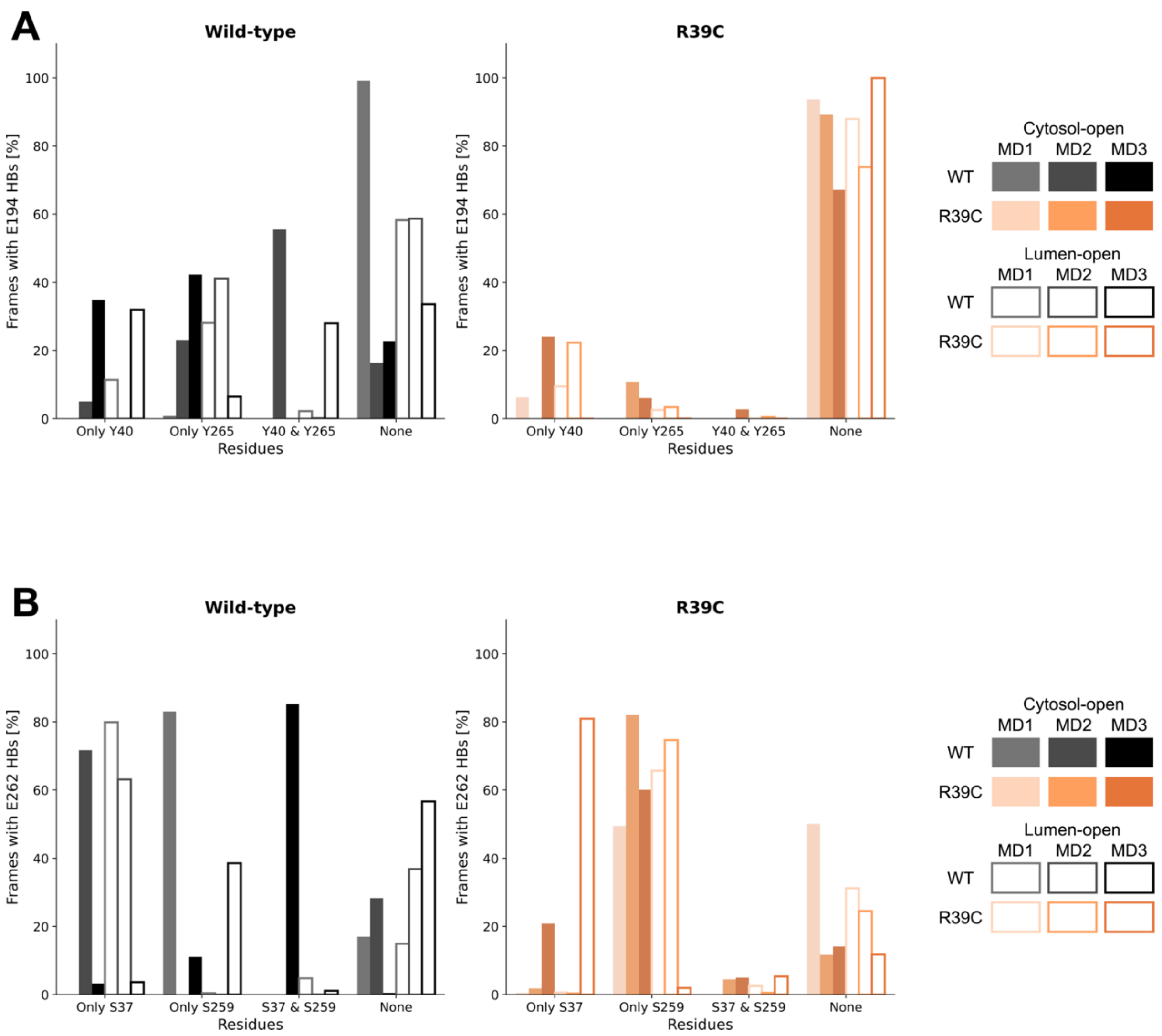


**Figure S15.** Hydrogen bonding interactions of E194 and E262. Barplots showing the percentage of frames in the MD simulations of the LO (unfilled bars) and CO (filled bars) conformations for wild-type (left) and R39C mutant (right) Sialin. A) Hydrogen bonding interactions between the E194 side chain carboxylate oxygens and the phenolic oxygens of Y40 alone, Y265 alone, both Y40 and Y265, or neither. B) Hydrogen bonding interactions between the E262 side chain carboxylate oxygens and the hydroxyl oxygens of S37 alone, S259 alone, both S37 and S259, or neither. The data shown are from the last 300 ns of the production run. Hydrogen bonds were determined to be present if the distance between the donor (D) and acceptor (A) was <3.5 Å and the angle A-D-H was <30°.

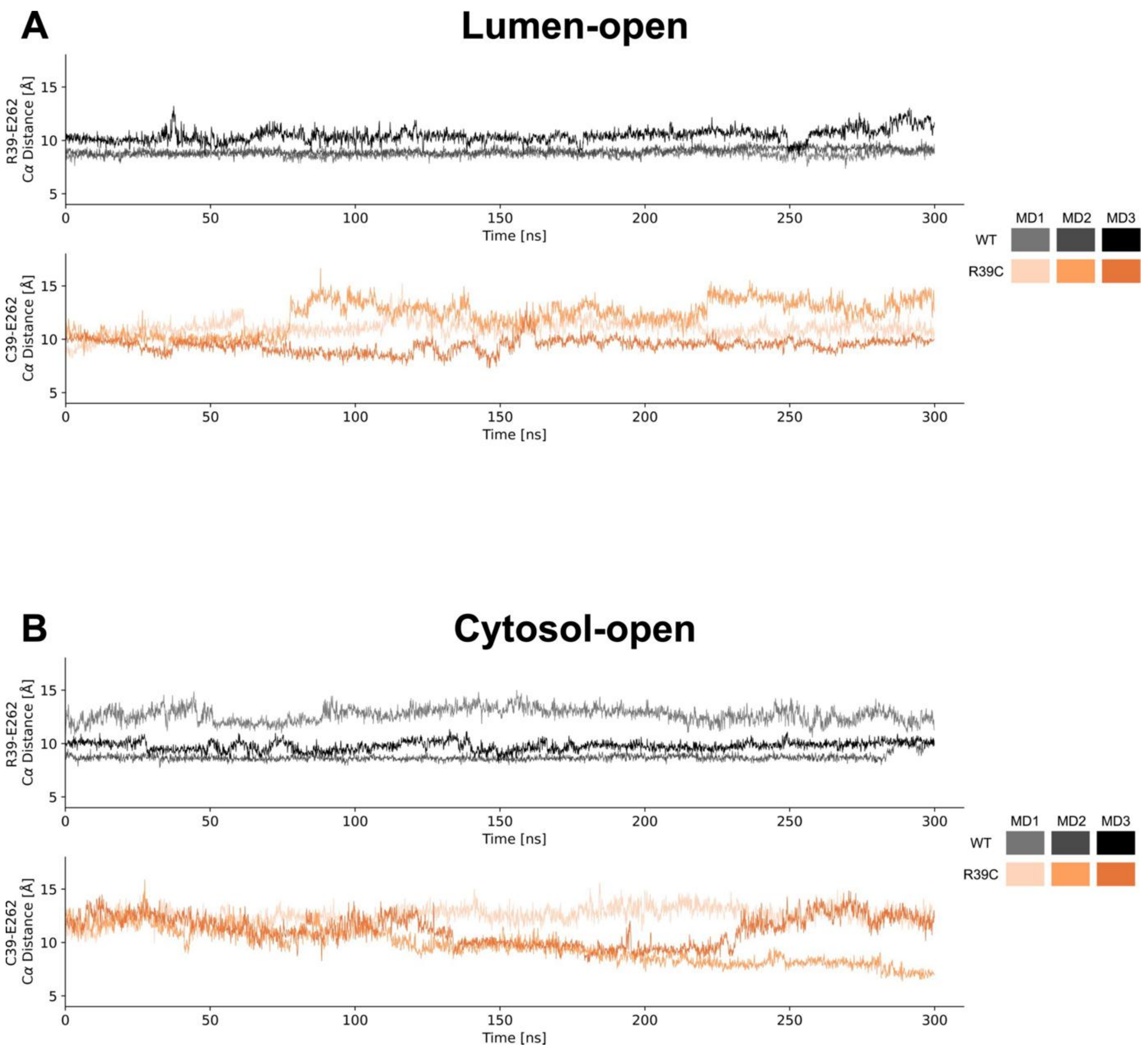


**Figure S16.** Line plots showing the evolution of the R39-E262 Cα distance. The evolution of the R39-E262 Cα distance in MD simulations of the LO (A) and CO (B) conformations is shown for wild-type (top, black) and R39C mutant (bottom, orange) sialin. The data of the last 300 ns of the production run of all three replicas, MD1 (lighter), MD2 (medium hue), and MD3 (darkest), are shown for each mutant.

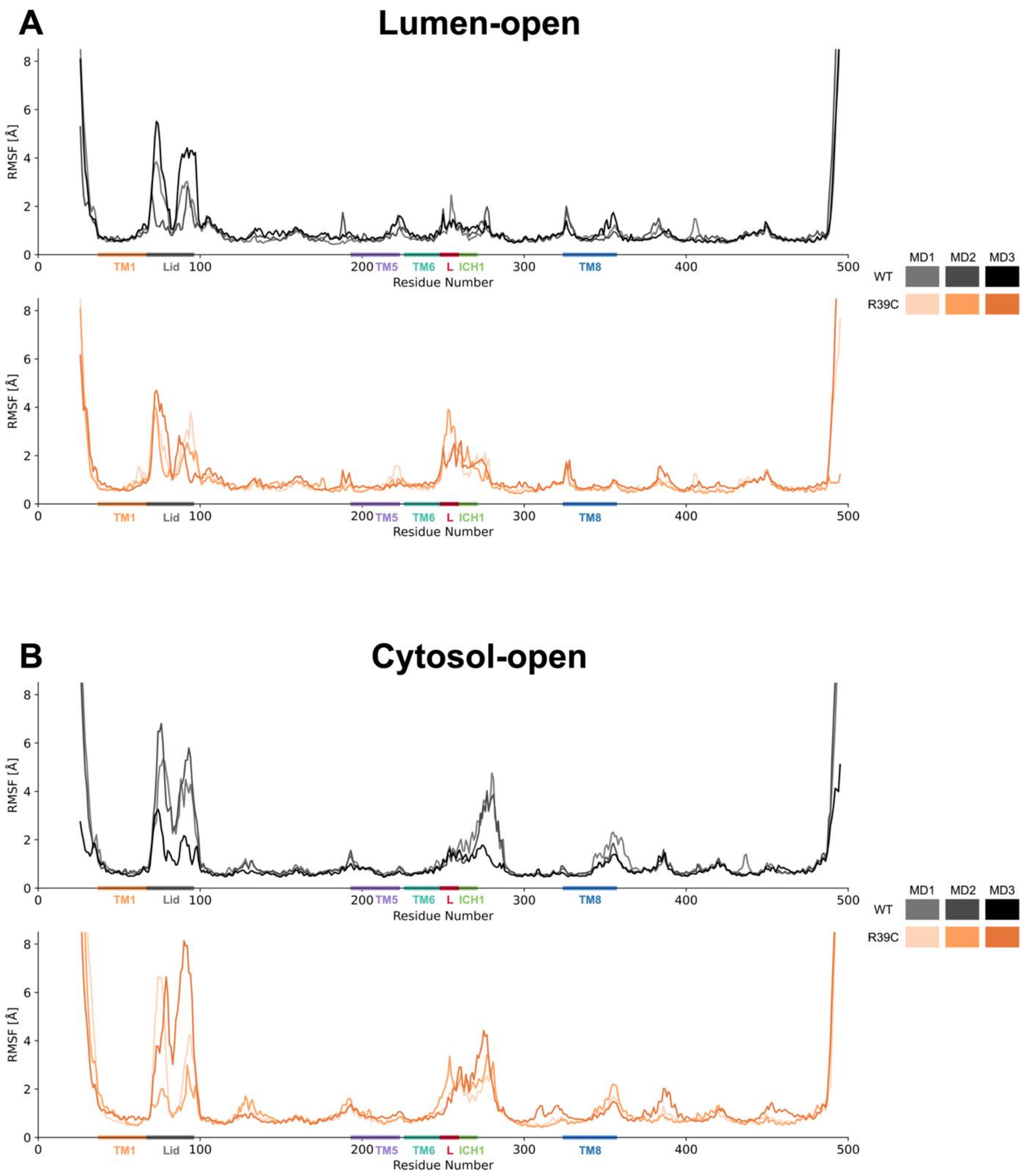


**Figure S17.** Line plots showing the RMSF values of the Cα atoms of the wild-type (top, black) and R39C (bottom, orange) mutant sialin during MD simulations of the LO (A) and CO (B) conformations. The TM1 (38-67, orange), lid (68-95, grey), TM5 (194-222, purple), TM6 (227-248, teal), TM6-ICH1 loop (249-260, L, red), ICH1 (261-270, green), and TM8 (325-356, blue) regions are highlighted. The data shown are from the three replicas: MD1 (lighter), MD2 (medium hue), and MD3 (darkest). The RMSF was calculated using the last 300 ns of the production run. RMSF values of N-and C-terminal residues that are higher than 8 Å are not shown.

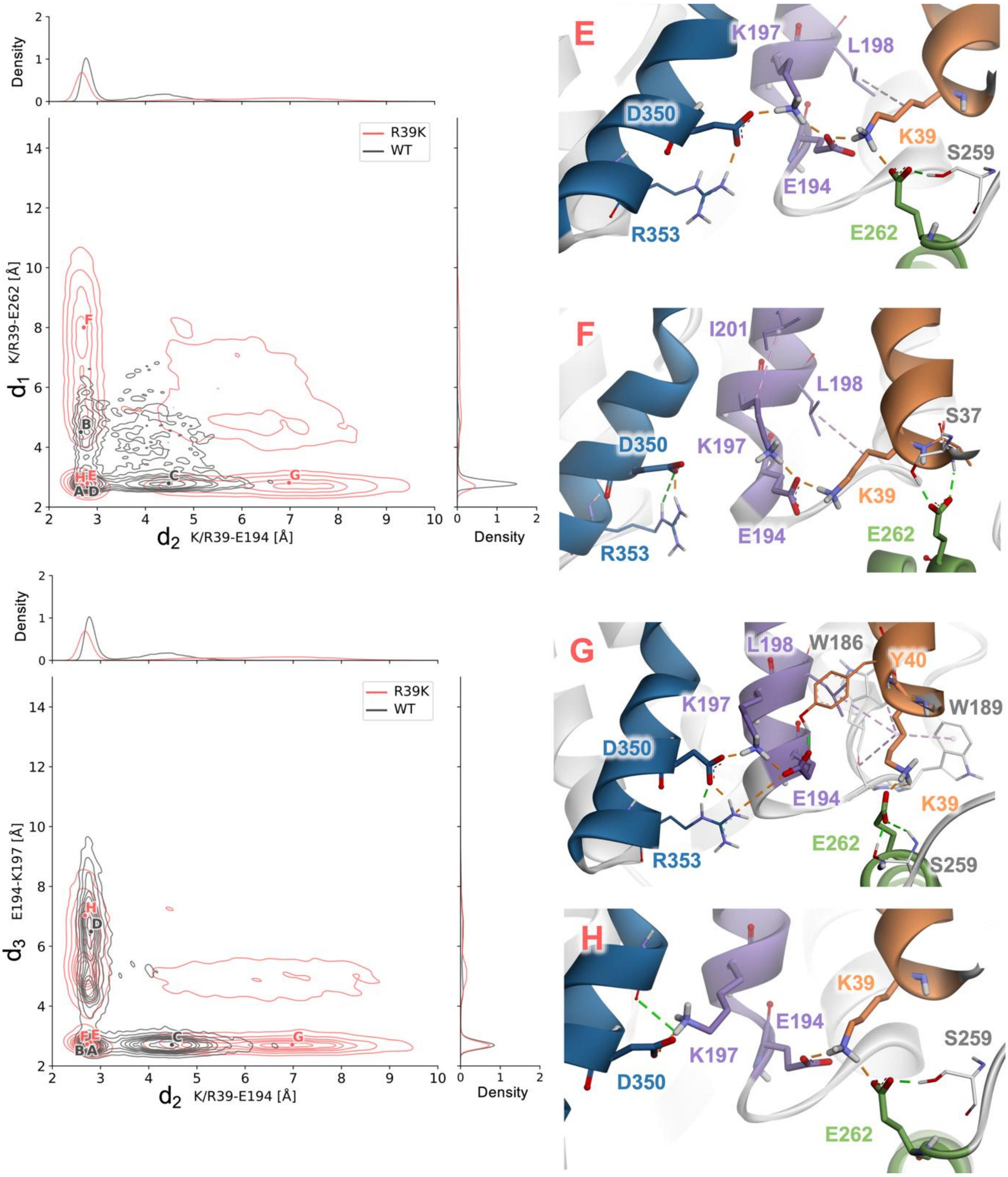


**Figure S18.** R39K mutant interactions in the LO conformation. Contour plots showing the minimum distances between the heavy atoms of the charged functional groups of K/R39, E194, K197, and E262. Plots show the K/R39-E194 ($d_2$) versus K/R39-E262 ($d_1$) (left) and K/R39-E194 ($d_2$) versus E194-K197 ($d_3$) (right) distances for the wild-type (black) and R39K mutant (red). Representative structures from the wild-type displayed in Figure S8 (A-D, black) and R39K mutant (E-H, red) are indicated with their corresponding distances highlighted in the contour plots. In wild-type sialin, the R39 site adopts four distinct configurations during the simulations displayed in Figure S8. In the R39K mutant of sialin, the interactions partially change: E) E194 forms strong interactions with both K39 and K197 and K39 simultaneously interacts strongly with E262; F) E194 maintains strong interactions with K39 and K197 and K39 moves away from E262 and does not remain within interaction range; G) E194 interacts strongly with K197 and is distant from K39 and not within interaction range, K39 forms a strong interaction with E262, and K197 interacts with D350; and H) K39, E194, and E262 form a strongly interacting triplet, E194 does not interact with K197, and K197 interacts with D350. The data from the last 300 ns of all three replicas (MD1-MD3) for both wild-type and mutant are shown. Residues in ICH1 (green) are shown to interact with TM1 (orange), TM5 (purple), TM8 (blue), and other parts (grey) of sialin. Hydrogen bonding (green dashed lines), charge-charge (orange dashed lines), and apolar (purple dashed lines) interactions are shown. Hydrogen bonds between backbone atoms involved in the formation of alpha-helices are not shown, and only polar hydrogens are displayed to improve visibility. Residues A26-C36 of the N-terminus are not shown to improve visibility.

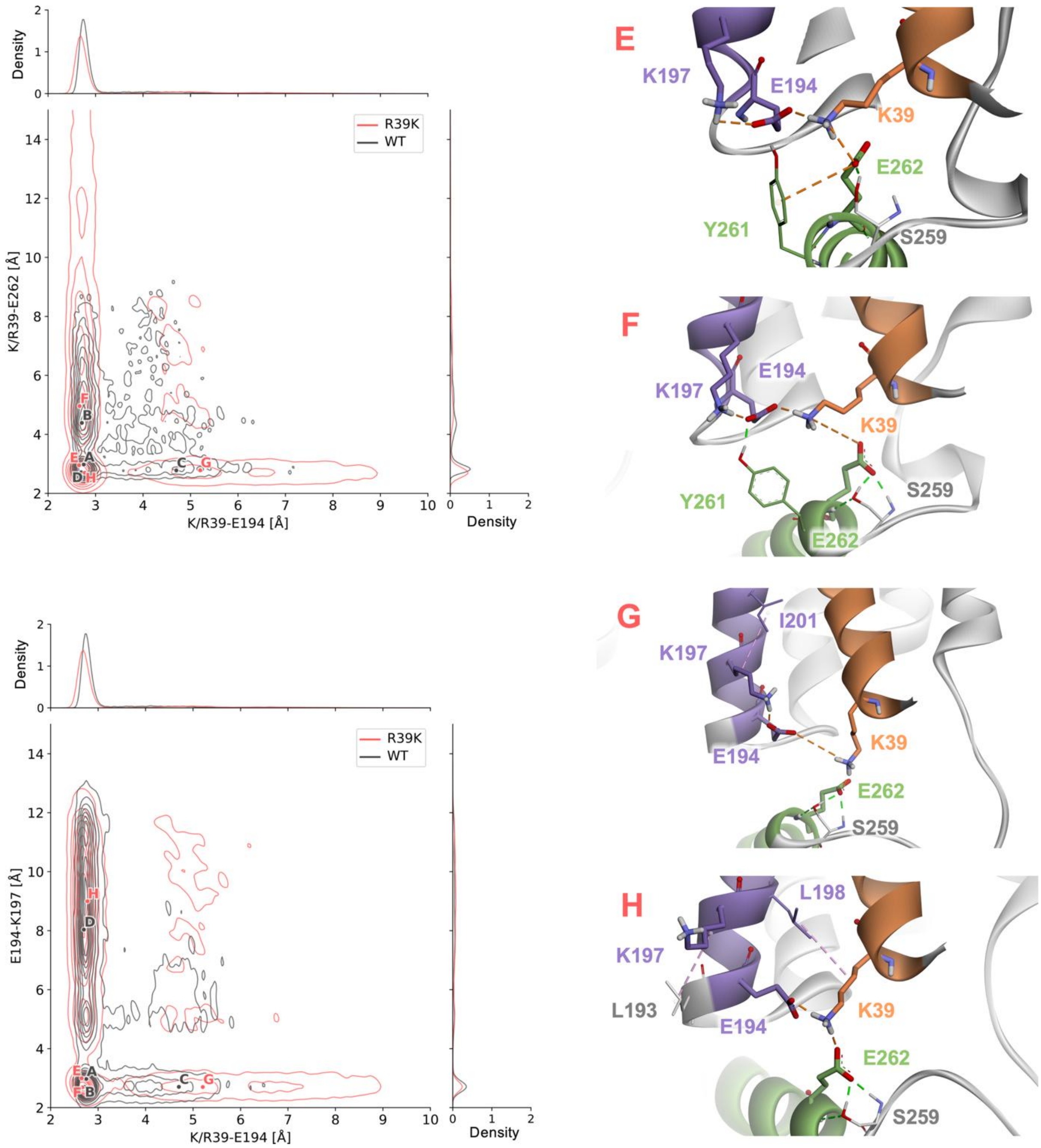


**Figure S19.** R39K mutant interactions in the CO conformation. Contour plots showing the minimum distances between the heavy atoms of the charged functional groups of K/R39, E194, K197, and E262. Plots show the K/R39-E194 versus K/R39-E262 (top) and K/R39-E194 versus E194-K197 (bottom) distances for the wild-type (black) and R39K mutant (red) sialin. Representative structures from R39K mutant (E-H, red) are shown with their corresponding distances highlighted in the contour plots, distances of the wild-type (A-D, black Figure S8) are reported on the plot. In the R39K mutant of sialin, the interactions partially change from WT: E) E194 forms strong interactions with both K39 and K197 and K39 simultaneously interacts strongly with E262; F) E194 maintains strong interactions with K39 and K197 and K39 moves away from E262; G) E194 interacts strongly with K197 and is distant from K39 and K39 forms a strong interaction with E262; and H) K39, E194, and E262 form a strongly interacting triplet and E194 does not interact with K197. The data from the last 300 ns of all three replicas (MD1-MD3) for both wild-type and mutant are shown. Residues in ICH1 (green) are shown to interact with TM1 (orange), TM5 (purple), TM8 (blue) and other parts (grey) of sialin. Hydrogen bonding (green dashed lines), charge-charge (orange dashed lines), and apolar (purple dashed lines) interactions are shown. Hydrogen bonds between backbone atoms involved in the formation of alpha-helices are not shown and only polar hydrogens are displayed to improve visibility. Residues A26-C36 of the N-terminus are not shown to improve visibility.

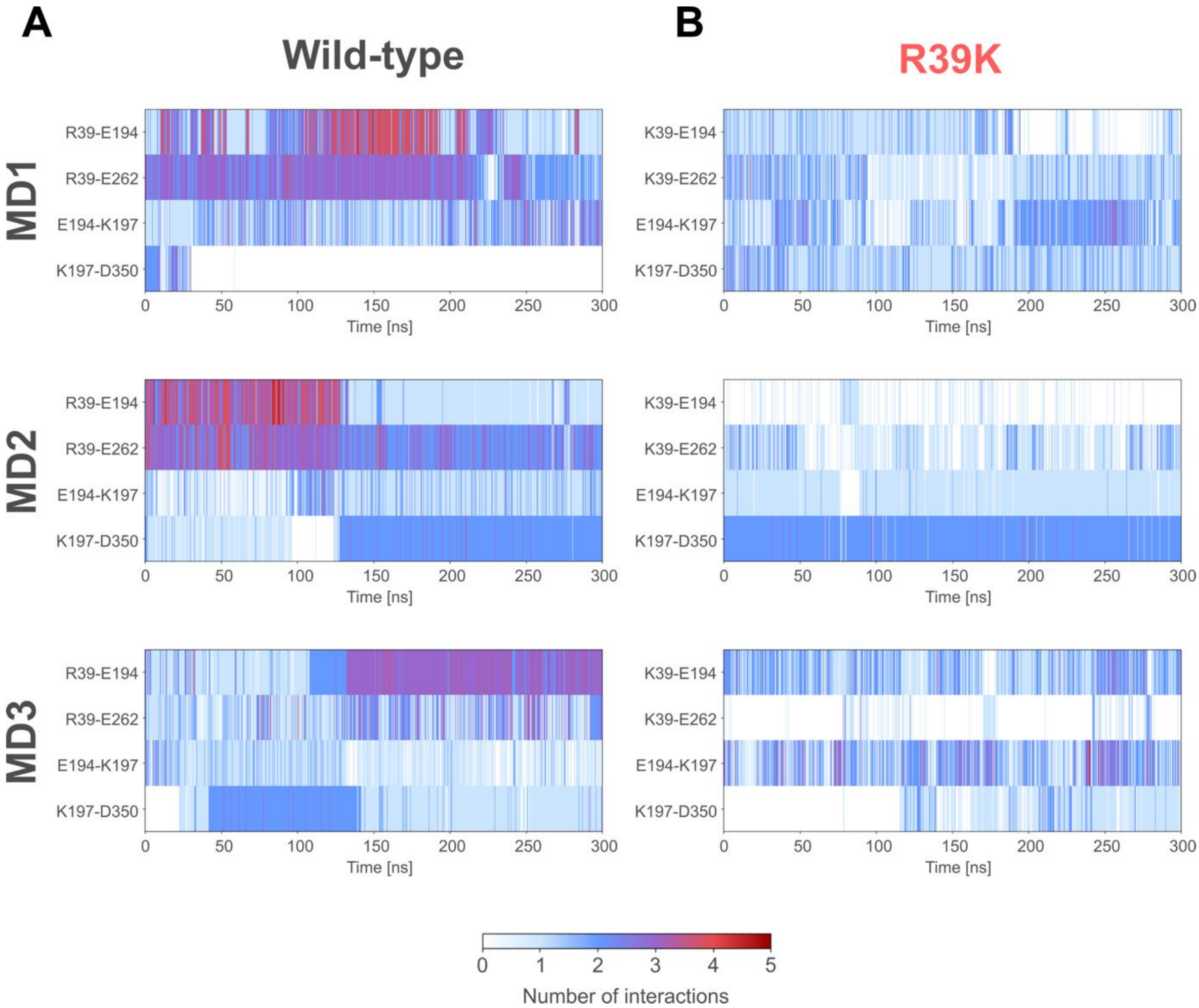


**Figure S20.** Non-bonded interaction monitoring of the wild-type and R39K LO trajectories. The interactions for the K/R39-E194, K/R39-E262, E194-K197, and K197-D350 pairs are shown during MD simulations for wild-type (left) and R39K mutant (right) sialin in the LO conformation. The data of the last 300 ns of the production run of all three replicas, MD1 (top), MD2 (middle), and MD3 (bottom), are shown for each mutant. Interactions are shown as vertical lines, where the colour of the line indicates the number of interactions, ranging from 0 (white) to 5 (red).

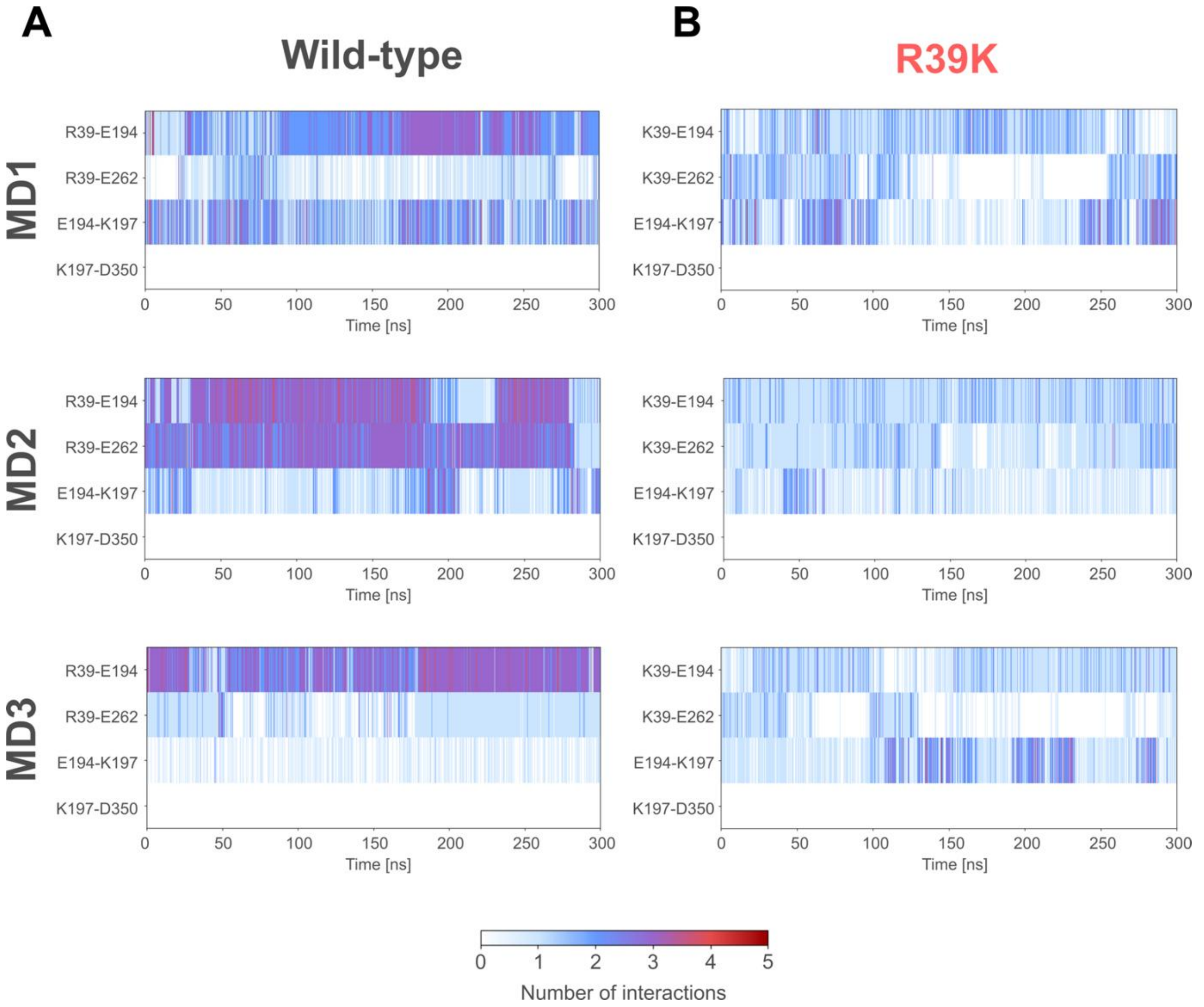


**Figure S21.** Non-bonded interaction monitoring of the wild-type and R39K CO trajectories. The interactions for the K/R39-E194, K/R39-E262, E194-K197, and K197-D350 pairs are shown during MD simulations for wild-type (left) and R39K mutant (right) sialin in the CO conformation. The data of the last 300 ns of the production run of all three replicas, MD1 (top), MD2 (middle), and MD3 (bottom), are shown for each mutant. Interactions are shown as vertical lines, where the colour of the line indicates the number of interactions, ranging from 0 (white) to 5 (red).

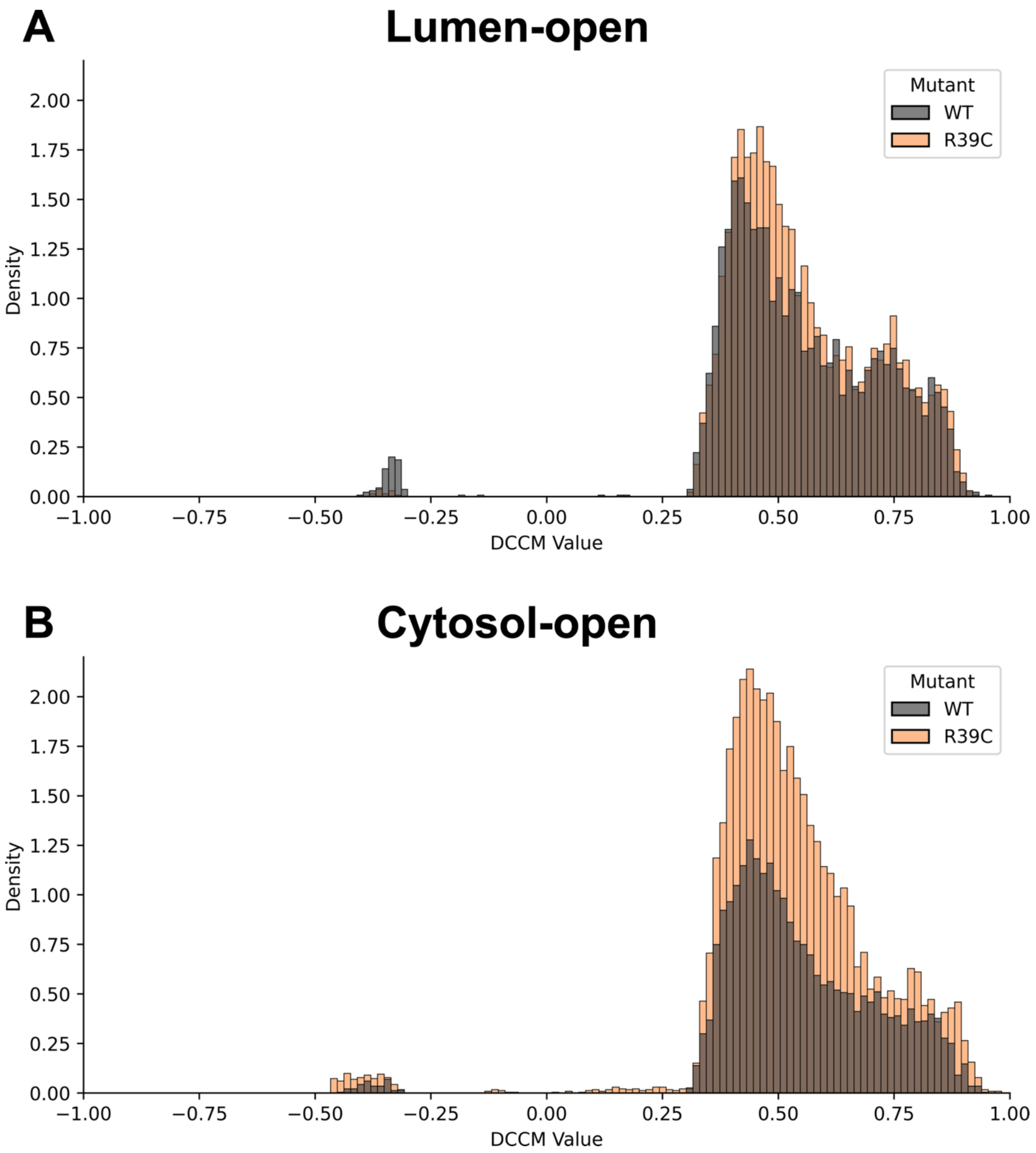


**Figure S22.** Histograms showing the distribution of dynamic cross-correlation (DCC) values between wild-type (WT) and R39C sialin mutant. The distributions of DCC values are shown for the LO (A) and CO (B) conformations of wild-type (WT, grey) and R39C mutant (orange) sialin. Only non-zero values in the lower triangle and not on the diagonal of the DCC matrix are shown.

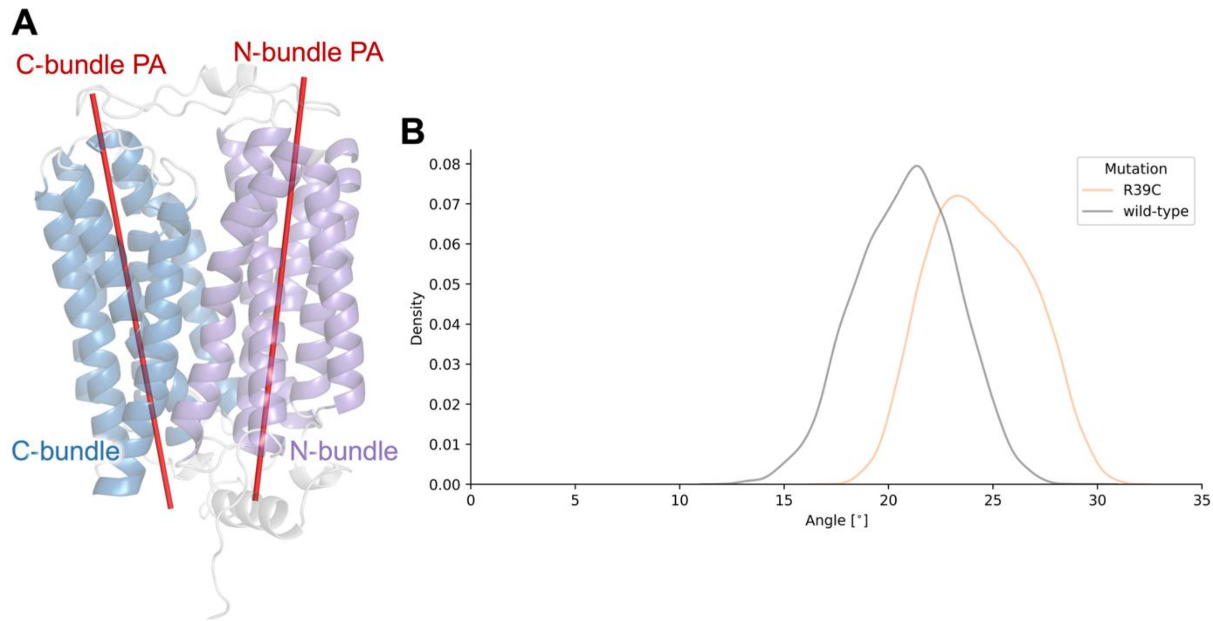


**Figure S23.** Bundle angle in the LO conformation of wild-type and R39C mutant sialin. A) The bundle angle was defined as the angle between the principal axes (PA, red) of the N-bundle (purple) and C-bundle (blue), derived from principal component analysis of the positions of the Cα atoms of the respective transmembrane helices. B) The distribution of the bundle angle in the wild-type (grey) and R39C mutant (orange) of sialin in the LO conformation are shown for the last 300 ns of the production run of all replicas (MD1- MD3) pooled together.

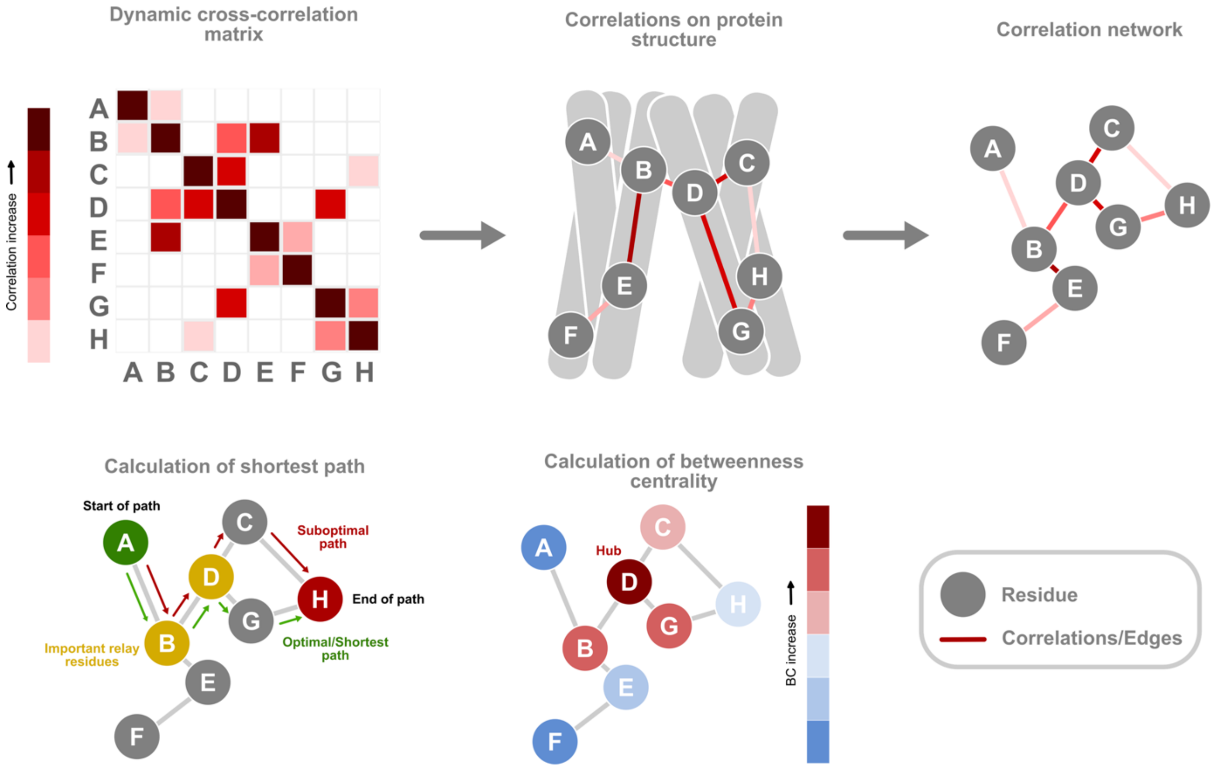


**Figure S24.** Schematic representation of network analysis. The dynamic cross-correlation matrix (top left) provides correlation values that indicate how coordinated the movement of two residues is. In the protein structure (top middle), residues (A-H) whose movement is correlated are connected over lines (edges) representing higher (darker red) and lower (lighter red) correlation coefficient values. The residues can be represented as nodes connected through edges in a correlation network (top right). In this network representation, the spatial arrangement of nodes reflects correlation strength rather than physical proximity in the protein structure; thus, residues distant in 3D space may appear close in the network if their movements are highly correlated. From this network, we calculate the shortest path connecting any two residues (e.g., from A to H, shown in green, bottom left), as well as alternative suboptimal paths (shown in red). Residues through which many optimal and suboptimal paths pass are important relay residues (yellow). By calculating the shortest path between every pair of residues, we can determine how many shortest paths pass through each residue. This value represents the betweenness centrality (bottom middle). Residues that are part of many shortest paths have high betweenness centrality and represent hubs of allosteric communication.

**Table S1** Primers used for site-directed mutagenesis. Mutated nucleotides are indicated in bold characters. Silent restriction sites (underlined) were introduced simultaneously to amino acid changes to facilitate screening of positive clones.

| **Mutations** | **Primers** | **Sens** | **Sequences (5' to 3')** |
|---|---|---|---|
| C36A | SIA_122S | Sens | TCCAGTGTGC**GC**CTCTGCTCGTTACAACTTAGCAATTTTGGCCTTTTTTG |
| | SIA_123A | Antisens | GCGGCTTCGGCCCGTGGG |
| S37A | SIA_124S | Sens | AGTGTGCTGC**GC**TGCTCGTTACAACTTAGC |
| | SIA_125A | Antisens | GGAGCGGCTTCGGCCCGT |
| S37C | SIA_126S | Sens | AGTGTGCTGCT**G**TGCTCGTTACAACTTAGC |
| | SIA_127A | Antisens | GGAGCGGCTTCGGCCCGT |
| A38R | SIA_150S | Sens | GTGCTGCTCT**CG**TCGTTACAACTTAGCAATTTTGGCC |
| | SIA_151A | Antisens | ACTGGAGCGGCTTCGGCC |
| A38R (R39C) | SIA_174S | Sens | GTGCTGCTCT**CGC**TGCTACAACTTAGCAATTTTG |
| | SIA_175A | Antisens | ACTGGAGCGGCTTCG |
| R39D | SIA_178S | Sens | CTGCTCTGCT**GA**TTACAACTTAGCAATTTTGG |
| | SIA_179A | Antisens | CACACTGGAGCGGCT |
| R39F | SIA_181S | Sens | CTGCTCTGCT**TT**TTACAACTTAGCAATTTTGG |
| | SIA_179A | Antisens | CACACTGGAGCGGCT |
| R39K | CS_SiaR39K_F | Sens | CTGCTCTGCT**AAG**TA**T**AACTTAGCAATTTTGGCCTTTTTTG |
| | CS_SiaR39K_R | Antisens | CACACTGGAGCGGCTTCG |
| R39M | SIA_128S | Sens | CTGCTCTGCT**ATG**TACAACTTAGCAATTTTGGCCTTTTTTGGTTTC |
| | SIA_129A | Antisens | CACACTGGAGCGGCTTCG |
| Y40A | SIA_118S | Sens | GCTGCTCTGC**A**CGT**GC**CAACTTAGCAATTTTGG |
| | SIA_119A | Antisens | ACACTGGAGCGGCTTCGG |
| E194A | SIA_114S | Sens | GCAA**G**CTTCTTAGCATTTCATATGCAGGAG |
| | SIA_115A | Antisens | TTCTT**G**CAAGAGGGGGAGCCCAAG |
| E194R | SIA_152S | Sens | TCCCCCTCTT**CG**AAGAAGCAAAC |
| | SIA_153A | Antisens | GCCCAAGAAGACCACATG |
| Y261A | SIA_156S | Sens | AATTTCCCAT**GC**TGAAAAGGAATACATTC |
| | SIA_157A | Antisens | CTCTTGTGTTTTTGTGGTG |
| E262A | SIA_116S | Sens | TCCCATTATG**C**AAAGGAATACATTC |
| | SIA_117A | Antisens | AATTCTCTTGTGTTTTTGTG |
| E262R | SIA_176S | Sens | TTCCCATTAT**CGC**AAGGAATACATTCTTTC |
| | SIA_177A | Antisens | ATTCTCTTGTGTTTTTGTGG |
| K263A | SIA_158S | Sens | CCATTATGAA**GC**GGAATACATTCTTTCATC |
| | SIA_159A | Antisens | GAAATTCTCTTGTGTTTTTGTG |
| E264A | SIA_164S | Sens | TATGAAAAGG**C**ATACATTCTTTCATC |
| | SIA_165S | Antisens | ATGGGAAATTCTCTTGTG |
| E264K | SIA_172S | Sens | TTATGAAAAG**A**AATACATTCTTTCATCATTAAG |
| | SIA_171A | Antisens | TGGGAAATTCTCTTGTGTTTTTG |
| E264P | SIA_132S | Sens | TTATGAAAAG**CC**ATACATTCTTTCATCATTAAG |
| | SIA_133A | Antisens | TGGGAAATTCTCTTGTGTTTTTG |
| Y265A | SIA_134S | Sens | TGAAAAGGAA**GC**CATTCTTTCATCATTAAGAAATC |
| | SIA_135A | Antisens | TAATGGGAAATTCTCTTGTG |
| I266A | SIA_120S | Sens | AAAGGAATAC**GC**TCTTTCATCATTAAGAAATCAG |
| | SIA_121A | Antisens | TCATAATGGGAAATTCTCTTG |
| L267A | SIA_195S | Sens | GGAATACATT**GCG**TCATCATTAAGAAATCAGCTTTCTTC |

| | | | |
|---|---|---|---|
| | SIA_196S | Antisens | TTTTCATAATGGGAAATTCTCTTGT |
| S268A | SIA_136S | Sens | ATACATTCTT**G**CATCATTAAGAAATC |
| | SIA_137A | Antisens | TCCTTTTCATAATGGGAAATTC |
| S269A | SIA_138S | Sens | CATTCTTTCA**G**CATTAAGAAATCAG |
| | SIA_139A | Antisens | TATTCCTTTTCATAATGGGAAATTC |
| L270A | SIA_193S | Sens | TCTTTCATCA**GCG**AGAAATCAGCTTTCTTC |
| | SIA_194A | Antisens | ATGTATTCCTTTTCATAATGG |
| L270S | SIA_183S | Sens | TCATCA**AGC**AGAAATCAGCTTTCTTC |
| | SIA_184A | Antisens | ATGTATTCCTTTTCATAATGG |

**Table S2.** Sequence of equilibration steps for the MD simulations.

| Step | Duration [ns] | Conditions | Restraints on protein heavy atoms [kJ/mol/nm2] | Restraints on lipid heavy atoms [kJ/mol/nm2] | Time step [fs] |
|---|---|---|---|---|---|
| 1 | 1 | NVT | 1000 | 1000 | 1 |
| 2 | 5 | NPT | 1000 | 1000 | 1 |
| 3 | 5 | NPT | 1000 | 500 | 1 |
| 4 | 5 | NPT | 1000 | 250 | 1 |
| 5 | 10 | NPT | 1000 | - | 1 |
| 6 | 5 | NPT | 500 | - | 1 |
| 7 | 5 | NPT | 500 (only Cα atoms) | - | 1 |
| 8 | 5 | NPT | 250 (only Cα atoms) | - | 1 |
| 9 | 20 | NPT | - | - | 1 |
| 10 | 30 | NPT | - | - | 2 |
